# Time-resolved brain network community detection based on instantaneous phase of fMRI data

**DOI:** 10.64898/2026.03.17.712372

**Authors:** Marika Strindberg, Peter Fransson

## Abstract

In this paper, we propose a novel method that estimates time-resolved communities, or networks, from parcellated fMRI data based on instantaneous phase of the parcel timeseries. Importantly, each community label reference a limited phase range across time and subjects. It thus avoids the relabelling problem that is common for community detection algorithms. Our aim was to enhance the temporal resolution of brain network analysis in a whole-brain context. The method provides insights into how brain regions synchronize both within and across subjects and reorganize during task performance. It offers a complementary perspective on network integration and segregation as concurrent processes, quantified through differences in instantaneous phase. We employed HCP motor task fMRI data to exemplify its practical application. Prior to calculation of instantaneous phase, signal timeseries were decomposed into two signal modes with minimally overlapping frequency ranges. We show that task specific motor movements (hand, feet, tongue) can be separated from block-design related activation (visual and attention networks) where the former was found in the slower mode and the latter in the higher frequency mode.

## 1. Introduction

Since the discovery of EEG a century ago by Hans Berger (Berger, 1929), interest in the dynamic activity of the brain has been ongoing (Quigley, 2022). With the invention of the bold-level-dependent (BOLD) fMRI signal in the 1990s (Ogawa et al., 1990), the precision in locating the anatomical origin increased significantly (Bandettini et al., 1992; Kwong et al., 1992). fMRI has since become an important cornerstone in the repertoire of brain investigation methods. Initially, fMRI studies mainly focused on using the improved spatial resolution to answer questions about which parts of the brain show increased activity in response to a particular task. A significant advancement in our understanding of the brain made possible by fMRI was the discovery of a functional large-scale organization of the brain into networks (Beckmann et al., 2005; Biswal et al., 1995; Bullmore & Sporns, 2009; Cordes et al., 2001; Damoiseaux et al., 2006; Fox et al., 2005; Friston, 1994; Greicius et al., 2003; Raichle et al., 2001; Smith et al., 2009; Sporns et al., 2000). A network typically consists of a set of distributed brain regions that systematically co-activate during a specified condition. The perhaps most well-known networks are the canonical intrinsic networks that are observed when a person is left awake in an MRI scanner without a specific task to solve (Smith et al., 2009; Yeo et al., 2011). Reflecting the internally focused “resting” condition, intrinsic networks are commonly referred to as resting-state networks (RSN). The highly organized nature and replicability of networks across subjects suggests that they are related to functionally meaningful “work” (Reid et al., 2019). However, because there are vastly more potential discrete “tasks” than large-scale networks in the brain, a more complex picture has gradually emerged in which each network is implicated in multiple tasks (Laird et al., 2009; Uddin et al., 2023; Yeo et al., 2015). This reflects a modular hierarchical structure, where networks consist of subnetworks that can be transiently engaged across tasks and become anti-correlated in relation to each other (Andrews-Hanna et al., 2010; Bassett et al., 2013; Chang & Glover, 2010; Cole et al., 2014; Fransson, 2005; He, 2013; Karahanoğlu & Van De Ville, 2015; Meunier, 2009; Uddin et al., 2009; Van De Ville & Liégeois, 2024; Vidaurre et al., 2017). Network activity is also situated in a whole-brain context characterized by a dynamic interplay between networks (He, 2013). A classic example is the deactivation of the default mode network (DMN) during externally focused task states (Fox et al., 2005; Fransson, 2005). Importantly, the state of the brain modulates network activity (Friston et al., 2000; Martin et al., 2021).

To better understand the rich repertoire of network engagement and interactions during conditions requiring both internally and externally focused attention, a number of methods for time-varying functional connectivity (TVFC) analysis have been developed, albeit not all of them take an explicit network perspective (Allen et al., 2014; Bassett et al., 2011a, 2013; Calhoun et al., 2014; Hutchison, Womelsdorf, Gati, et al., 2013; S. Keilholz et al., 2017; Lurie et al., 2020; Mucha et al., 2010; Preti et al., 2017; Thompson et al., 2018; Thompson & Fransson, 2018; Vidaurre, 2024). They can be broadly grouped into three main categories: (1) Computational mechanistic models based on theoretical frameworks of how brain signals are generated, typically derived from dynamical systems theory (Breakspear, 2017) (e.g., dynamic mean field models (Deco et al., 2013; Wong & Wang, 2006)). Real data is fitted to the model, and the accuracy of the model predictions are evaluated. (2) Statistical models that define the signal dynamics mathematically and fit the data to the model (e.g., state-space models such as Hidden Markow-models (HHM) (Vidaurre et al., 2017), Dynamical Conditional correlation (DCC) (Lindquist et al., 2014), Dynamic Community Detection (Bassett et al., 2013)). (3) Data-driven methods that identify patterns from the data directly without a prior model (Lurie et al., 2020) (e.g. time-frequency analysis (Yaesoubi et al., 2015)(Chang & Glover, 2010), Co-activation patterns (CAP) (Liu et al., 2018), innovation-driven co-activation patterns (iCAP) (Karahanoğlu & Van De Ville, 2015), sliding-window correlation (Allen et al., 2014; Handwerker et al., 2012; Leonardi et al., 2013; Sakoğlu et al., 2010; Tagliazucchi et al., 2012), Leading Eigenvector Dynamics Analysis (LEiDA) (Cabral et al., 2017), and instantaneous phase synchrony analysis (IPSA) (Cabral et al., 2017; Glerean et al., 2012; Honari et al., 2021; Niazy et al., 2011; Pedersen et al., 2018; Ponce-Alvarez et al., 2015; Preti et al., 2015; Strindberg et al., 2021)). For details and complete lists of methods see review papers (Hutchison, Womelsdorf, Allen, et al., 2013; S. Keilholz et al., 2017; Lurie et al., 2020; Preti et al., 2017).

In this study, we present a novel data-driven approach within the IPSA group, aimed at enhancing the temporal resolution of brain network analysis in a whole-brain context. Our proposed method estimates time-resolved clusters, or networks, based on the instantaneous phase of the fMRI signal. This method respects the temporal order (Lurie et al., 2020) of the data and captures changes in whole-brain dynamics from one time point to the next. Importantly, community labels refer to the same property, i.e. a limited phase distribution, across time and subjects and thus avoids the relabelling problem that is common for community detection algorithms (Khambhati et al., 2018). The method offers a complementary perspective on network integration and segregation as concurrent processes, quantified through differences in the instantaneous phase.

The Human Connectome Project (HCP) Motor task data (Glasser et al., 2016; Uğurbil et al., 2013) was utilized to illustrate the effectiveness of the method. For clarity, in this methodological paper, we focus on the general principles of the method and its applicability to group-level analysis. We will refrain from in-depth analysis and discussion of all relevant patterns in the dynamic brain network activity revealed by the method. This discussion, together with a presentation of additional questions that can be addressed with the method, such as the temporal order of network recruitment within subjects and on the group level, will be presented in a separate paper.

## 2. Methods

### 2.1 fMRI Data

fMRI data from the HCP Young Adult 900 subject re-processed release (August 2025) were included in this study (Glasser et al., 2016; Uğurbil et al., 2013; Van Essen et al., 2013). Data collection was approved by Washington University institutional review board. Informed consent was provided by subjects. 791 subjects with complete 3T task data were considered for inclusion. The two motor fMRI task runs (block design, scanned back-to-back) used here were acquired at the end of a close to a one-hour fMRI recording session (Barch et al., 2013). In the first run (run A), phase encoding was in the right-left direction, and in the second run (run B), phase encoding was in the left-right direction. Subjects with head movement with a peak frame-wise displacement root mean square (fdrms) > 0.5 mm were excluded from further analysis. After considering the degree of subject head motion, 512 subjects remained for run A, and 518 subjects remained for run B. We present results from run B. The motor task consisted of five types of movements (finger-tapping left hand, finger-tapping right hand, toe squeezing right foot, toe squeezing left foot, and tongue movement)(Barch et al., 2013). The duration of each block was 12 s and was preceded by a 3 s cue. Each type of movement/task block was repeated once. Additionally, three fixation blocks (duration: 15 s) were interspersed between the motor task blocks.

### 2.2 Preprocessing and parcellation

Minimally pre-processed volume-based data provided by the HCP were used and no further pre-processing of the data was conducted (Glasser et al., 2013). The pre-processing pipeline included gradient unwarping, Spin-echo ("SEBASED") intensity bias field correction, nonlinear registration to the MNI reference space using the FSL tool FNIRT, intensity normalization and multi-run FIX denoising (Barch et al., 2013; Glasser et al., 2013).

For the cortical parcellation, the Schaefer 400 brain area parcellation (seven networks) was used (Schaefer et al., 2018). The subcortical section of the Human Brainnetome Atlas (Fan et al., 2016) was added to delineate the thalamus, basal ganglia, hippocampus, and amygdala. In total, the parcellation comprised of 436 brain parcels. The amygdala and hippocampus parcels were assigned to the “limbic network” in the Schaefer parcellation scheme, whereas the thalamus and basal ganglia were treated as separate networks. The analysis therefore considered nine separate networks: Visual (VIS), Somato-motor (SOM), Dorsal attention (DAN), Ventral attention (VAN), Limbic (Limb), Frontoparietal (FPN), Default Mode (DMN), Thalamus (Thal), Basal Ganglia (BG). This was the same parcellation principle used in previous studies (Fransson & Strindberg, 2023; Strindberg et al., 2021).

### 2.3 Phase synchronization and instantaneous phase

A real valued signal S can be represented in its analytical form (i.e. complex valued), *S* = *A* × *e^iθ^* = *A*(cos(*θ*) + *i* sin(*θ*)), where A is the amplitude of the signal, θ is the phase angle and *i* the imaginary unit. In polar coordinates, the cosine term represents the real part of the phase angle, and the sine component represents its imaginary part. Phase synchronization between two different signals is defined as the degree of alignment of their respective phases. Phase synchronization is a classic measures in MEG and EEG studies, and increasingly so for fMRI data analysis (Cabral et al., 2017; Fransson & Strindberg, 2023; Glerean et al., 2012; Honari et al., 2021; Niazy et al., 2011; Ponce-Alvarez et al., 2015, 2015; Strindberg et al., 2021), to assess the degree of co-activation between regions of the brain (Glerean et al., 2012; Varela et al., 2001). The underlying assumption is that a small difference in phase translates into increased information exchange between regions (Varela et al., 2001). In analogy with anti-correlation, two timeseries with opposite phases can be said to be in anti-phase synchronization. Importantly, signal phase synchronization can also be computed for each signal intensity timepoint by estimating the instantaneous phase using the Hilbert transformation. An important methodological strength of phase as a measure of interregional relationships in the brain, is that it quantifies relationships on a continuous spectrum of positive and negative values (Cabral et al., 2017; Glerean et al., 2012; Honari et al., 2021; Ponce-Alvarez et al., 2015; Preti et al., 2015; Strindberg et al., 2021). This is conceptually similar to correlation, which spans positive and negative dimensions bounded by -1 to 1. Phase values can be made to be bounded in the same way by considering only the cosine of the phase (cos (*θ*)) or the sine sin (*θ*)(Cabral et al., 2017; Fransson & Strindberg, 2023; Glerean et al., 2012; Honari et al., 2021; Ponce-Alvarez et al., 2015; Strindberg et al., 2021). Sliding-window correlation and instantaneous phase measures have shown to yield similar results in estimating time-varying connectivity (Honari et al., 2021; Hutchison, Womelsdorf, Allen, et al., 2013; S. Keilholz et al., 2017; Pedersen et al., 2018; Preti et al., 2017).

### 2.4 Variational Mode Decomposition (VMD) of fMRI motor task data

The instantaneous phase extracted using Hilbert transform must be unambiguous. Thus, riding waves, which are typically present in real-world signals, must be eliminated. Narrow-band pass filtering is the most common method for making a time series smooth with unambiguous phase representation at each time point (Zatman, 1997). Typically, 0.01-0.1 Hz is the frequency range of the BOLD-signal that is considered coupled to neuronal activity (Biswal et al., 1995; S. D. Keilholz, 2014). However, band-pass filtering in this range will not eliminate riding waves and will therefore not constitute narrow band-pass filtering. However, which band-with to choose is not straight forward since it remains an open question which BOLD-signal frequency contain neuronally associated information(He et al., 2008). It has been demonstrated that the classic range can be divided into subranges typically referred to as slow-5 and slow-4 relating to neurophysiological counterparts (Zuo et al., 2010). There might also a wider range of meaningful frequencies within the BOLD-spectrum (Niazy et al., 2011). There seems to be task-specific segregation of network activation in different frequency bands, which is also present to some extent during rest (Billings et al., 2017; Chang & Glover, 2010; He et al., 2010; Thompson & Fransson, 2015; Yaesoubi et al., 2015). Thus, a single narrow filter is likely to result in information loss, which may be relevant. In addition, the choice of frequency cutoff is often arbitrary.

Signal mode decomposition is an alternative to narrow-bandpass filtering for extracting the instantaneous phase in fMRI data (Honari & Lindquist, 2022; Niazy et al., 2011). The empirical mode decomposition (EMD) algorithm was the first decomposition method proposed to divide a time series into intrinsic mode functions (IMFs), each with its own frequency profile, which together represent the original signal (Huang et al., 1998). The EMD algorithm is heuristic and iterative and thus sensitive to noise, which can result in significant mode mixing that occurs either with large amplitudes or when the frequencies constituting the original signal are too close to each other (Dragomiretskiy & Zosso, 2014; Xu et al., 2019). Several additional mode decomposition algorithms have been developed to mitigate the weaknesses of the EMD, some of which have been applied to fMRI data (for an overview, see Honari et al. 2022) (Honari & Lindquist, 2022). One of these methods is the variational mode decomposition (VMD) (Dragomiretskiy & Zosso, 2014). Within the VMD method, the number of modes is chosen a priori. The algorithm optimizes the decomposition such that modes with their respective frequency content reconstruct the original signal. In contrast to the EMD, the VMD algorithm is non-iterative. Instead, all modes are computed simultaneously. Compared to the original empirical mode decomposition (EMD) algorithm and the noise-extended Complete Ensemble Empirical Mode Decomposition (CEEMD) (Torres et al., 2011), the VMD has an improved mode separation and does not create artificial negative deflections prior to a strong positive deflection, as is seen using (CE)EMD). Considering these factors, we chose the VMD method to extract the modes. To further improve mode separation, data were first bandpass-filtered [0.01-0.15 Hz]. The lowpass threshold was chosen so that breathing artefacts from slow breathers (10 breaths/min corresponding to 0.17 Hz) would not be included. We also chose a prior that the VMD algorithm should extract two IMFs. This choice was motivated by our prior results using CEEMD on the same dataset. Notably, the CEEMD estimates the number of IMFs directly from the data through a heuristic sifting process, indicating the “true” number of modes in the data. For simplicity, in the remainder of this report, we will refer to the IMFs as “modes”.

Owing to edge effects from the band-pass filtering and VMD, 10 time points at the beginning and end of each fMRI signal time series were removed.

### 2.5 Phase clustering algorithm

As a starting point, we defined the dense weighted matrix *M_t_* where *t* denotes time. Dense means that all pairwise relationships between parcels were included without thresholding. Each matrix element *M_t_*(*x*, *y*) is an edge that describes the cosine of instantaneous phase difference *θ_x,y_* between parcels (nodes) *x* and *y* at time *t*. This algorithm is greedy and has a hierarchical structure. It optimizes two main criteria: integration (criterion one) and segregation (criterion two). Integration in this context means that *all* pairwise relationships within the same community have a maximum phase difference defined by threshold one, *thr*1 = cos (*θ*_1_). We investigated six different choices for 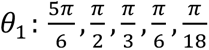. A schematic of the phase clustering algorithm is shown in Figure 1.

**Figure 1.**
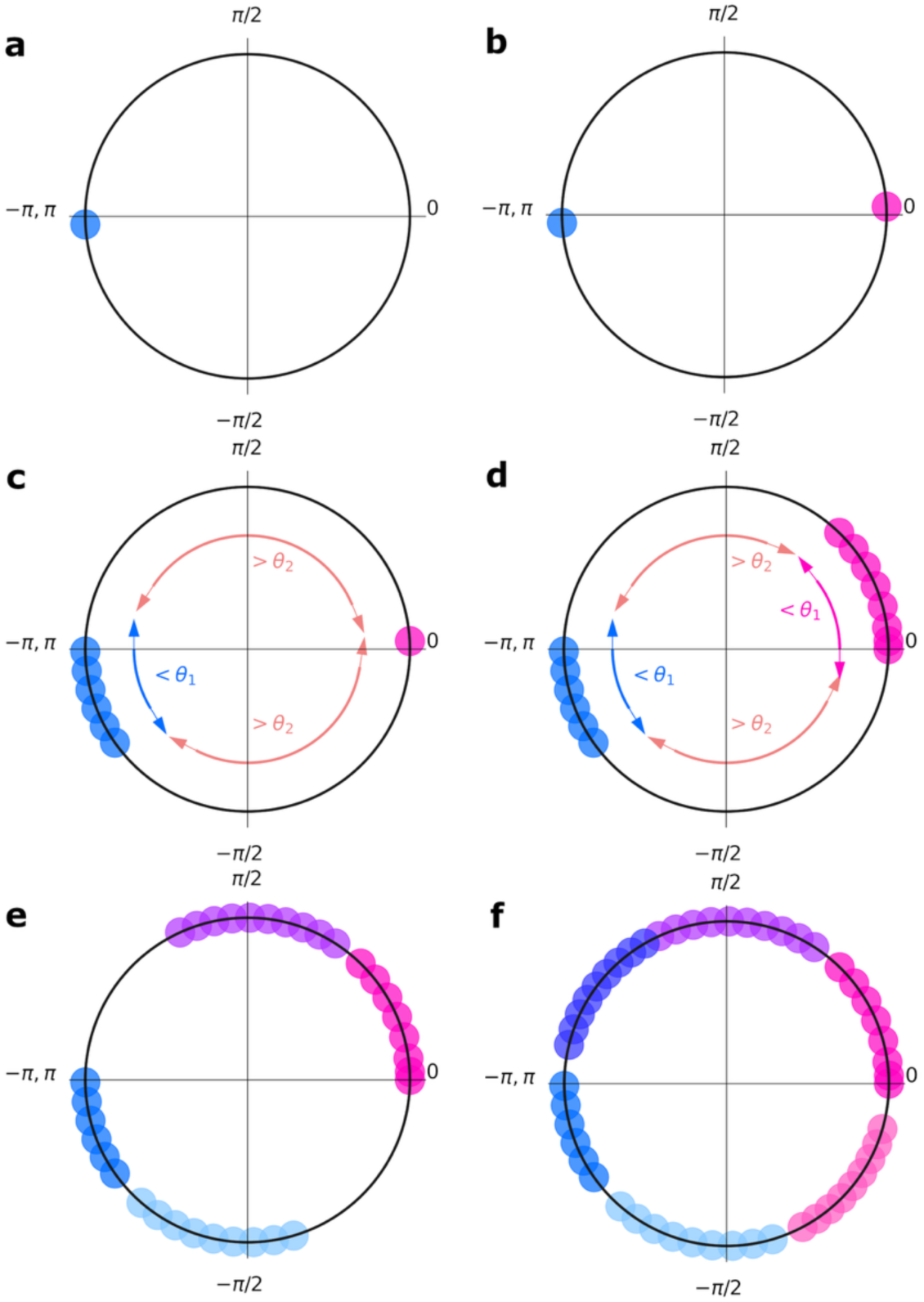
Schematic illustration of the principal steps in the phase cluster algorithm. At each timepoint, it assigns all parcels into communities based on two thresholds, integration threshold *thr*1 = cos (*θ*_1_) and segregation threshold *thr*2 = cos (*θ*_2_), where *θ*_2_ = −*θ*_1_. These relationships reflect the instantaneous phase difference between any two parcels within a community (thr1) and between parcels in the maximally segregated communities (thr2). In this example we used 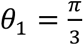 and *θ*_2_ = −*θ*_1_ which typically produces six communities. This means that all pairwise relationships (instantaneous phase) within a community are always < *θ*_1_ and all pairwise relationships between maximally segregated community are always > *θ*_2_. (a) For reproducibility purposes, the seed parcel (blue dot) for the first community is always selected to be the parcel with smallest instantaneous phase value. (b) The most segregated parcel in relation to the seed parcel in (a) is selected as the seed parcel for the second community. (c) First community is expanded according to thr1 and thr2. (d) Second community is expanded. Results after the first iteration. (e) Second iteration completed, that is, in total, four communities. (f) Third iteration completed; all parcels are assigned to a community, that is, six communities in total in this example.

**Figure 2.**
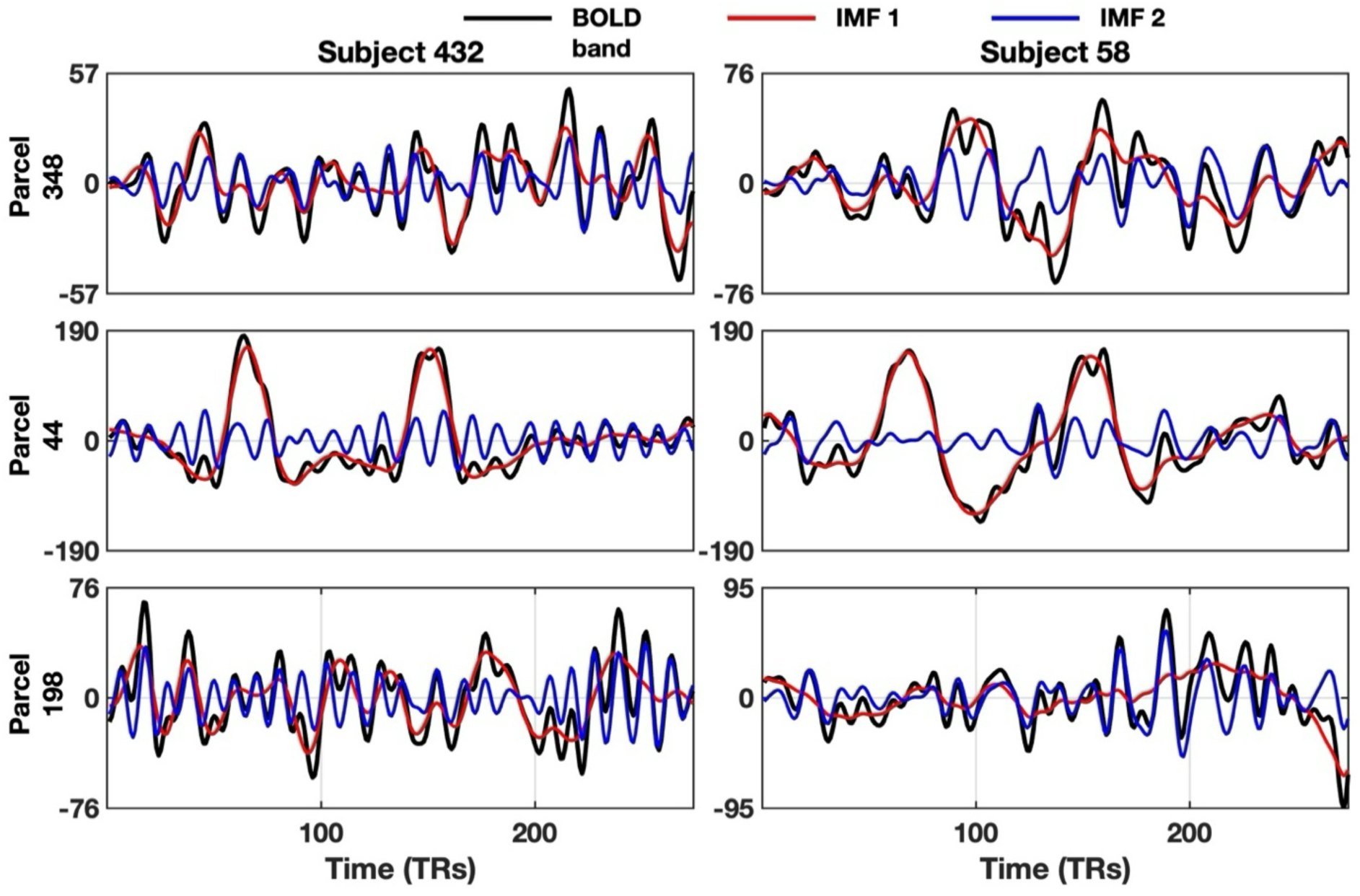
Example of VMD decomposition of motor task fMRI data. fMRI time-series were drawn from two randomly selected subjects (left and right columns, respectively) and from three parcels (rows). The band-pass filtered BOLD signal is shown in black, and the lower (mode 1) and higher (mode 2) frequency band VMD decomposed signal modes are shown in red and blue, respectively. Time series are normalized, and the y-axis is in arbitrary units.

In this context, an integrated community of parcels refers to a community that exhibits *phase consistency*, meaning that the maximum phase difference between any pair of parcels within the community is > *thr*1 = cos(*θ*_’_) (Recall that small values of *θ*_1_ translates to large *thr1)*. Segregated means that *all* pairwise relationships between nodes in two different communities have a *minimum phase difference* defined by threshold two, *thr*2 = cos (*θ*_2_), where *θ*_2_ = −*θ*_1_ if thr1 > 0 and otherwise *θ*_2_ = *θ*_1_. For each iteration, the algorithm aims to identify a pair of maximally segregated (criterion two), phase-consistent (integrated) communities (criterion one) if a minimum segregation of *at least thr*2 exists in the data. In each iteration, to harmonize communities of parcels across time points, the first community is based on the parcel with the smallest instantaneous phase value among the unassigned parcels, that is - *π* in the idealized case (see Fig. 1a). The second community is defined based on the parcel that is most segregated from the seed parcel of the first community (Fig. 1b). The first community is expanded by finding groups of parcels that fulfil both criteria one and two (Fig. 1c). If multiple such groups exist, the group with the smallest mean phase difference from the seed parcel is selected. The second community is expanded in the same manner (Fig. 1d). Communities are identified in the same way in subsequent iterations until all parcels were assigned to a community (Figs. 1e-f). A permutation option is available that optimizes for the largest of community in the expansion step of the communities after seed selection (Fig. 1c, d).

The results for 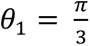 are presented in the main text (permutations not used). The algorithm is designed for community comparison between multiple iterations of a network, where the distribution of phases approximately span [−*π*, *π*]. The algorithm can be adapted to situations in which the phase distribution varies widely between the iterations and do not span [−*π*, *π*]. It could also be used with at different thr2 or without thr2.

### 2.6 Cross-subject phase synchronization

The phase clustering algorithm organizes parcels into phase-consistent communities at each temporal point within fMRI data runs obtained from individual subjects. As communities are designed to span a limited phase range defined by *thr1* and *thr2*, communities with the same label are comparable in terms of instantaneous phase content across time and subjects. For each mode, this provides a foundation for evaluating the similarity in brain responses within an individual at different temporal points. The method also facilitates the estimation the degree of synchronization across subjects for each parcel and mode. We defined the degree of cross-subject phase synchronization for each parcel, mode, and time point as the percentage of subjects across the entire cohort with the given parcel assigned to a particular community. The degree of cross-subject synchronization was computed separately for each community. Thus, the method provides a measure of the extent to which the phase of each parcel timeseries synchronized across subjects, on a time-point by time-point basis.

### 2.7 Dynamic estimation of network integration and segregation

By the thresholding criteria of the clustering procedure given above, communities computed with our method correspond to well-defined and quantifiable instantaneous phase relationships between parcels. Parcels in the same community are phase consistent (see above) and have clearly defined relationships with parcels in other communities. The relationships between communities exists on a gradient of integration and segregation. The highest level of integration is observed within communities. The level of segregation between communities depends on their relative positions to each other. Neighbouring communities are less segregated (in fact, certain subgroups of parcels in neighbouring communities might be integrated) than two communities on the opposite end of the unit circle (Fig. 1). The relative percentage of parcels (either in relation to the total number of parcels in parcellation or a to a specific network, depending on the research question) within communities can thus be used as a dynamic measure to estimate the average (across subjects) degree of integration and segregation at a time point by time point basis during an fMRI task run. At least three conceptually distinct types of integration-segregation scenarios are possible: a) a large fraction of parcels is affiliated with either of two maximally segregated communities (e.g., communities 1 and 4, 2 and 5, and 3 and 6, respectively); b) parcels are approximately equally scattered between all communities; and c) a large fraction of parcels is found in one dominating community, and the remainder of parcels are scattered among the other communities. We refer to these scenarios as “bimodal segregation,” “equalized”, and “single dominating” (see Suppl. Fig. S1 for an illustrative example).

### 2.8 Phase-shuffled data

To test the alternative hypothesis that any synchronization was driven by random processes, the algorithm was applied to surrogate data. Because the key property of the presented method is phase, phase shuffled data were used as surrogates (Cordes et al., n.d.; Lancaster et al., 2018; Miller et al., 2018). A shuffling of the phases preserves the power spectrum of a time course while destroying the temporal order of the phase (Suppl. Fig. S2). The shuffled results were used as a reference from which the statistical thresholds for significance were extracted. For both modes, results beyond ± 3*SD* were considered significant (Suppl. Figs. S3a, b).

## 3. Results

### 3.1 VMD-decomposition of fMRI motor data

The two extracted VMD modes had distinct power spectra with minor overlap (Suppl. Fig. S4). The first mode (lower frequencies) had two peaks at 0.01578 Hz and 0.03157 Hz, reflecting a time-period of 63.4s and 31.8 s respectively. The second mode (higher frequencies) had a single distinct peak at 0.06839 Hz (i.e. a time-period of 15.7 s). The latter closely corresponds to the block design of 15 seconds consisting of a 3 s cue and 12 s task-blocks or 15s fixation-blocks. Excluding the SOM (Somato-motor canonical resting-state network) parcels substantially diminishes the two peaks in mode 1 but not mode 2. No such effect was seen when a random selection of the same number of parcels (n = 77) was removed (Suppl. Fig. S5a). Similarly, the major peak in mode 2 was largely reflected by the activity of the visual (VIS) and the VAN and DAN (ventral and dorsal attention networks, respectively) (Suppl. Fig. S5b).

### 3.2 Cluster results

The choice of integration/segregation thresholds (thr1 = cos (*θ*_1_), thr2 = cos(*θ*_2_) *w*ℎ*ere θ*_2_ = −*θ*_1_) directly effects the number and sizes of communities. This is illustrated in Fig 3 for the lower frequency mode (mode 1); however, the results were similar for the higher frequency mode (mode 2).

**Figure 3.**
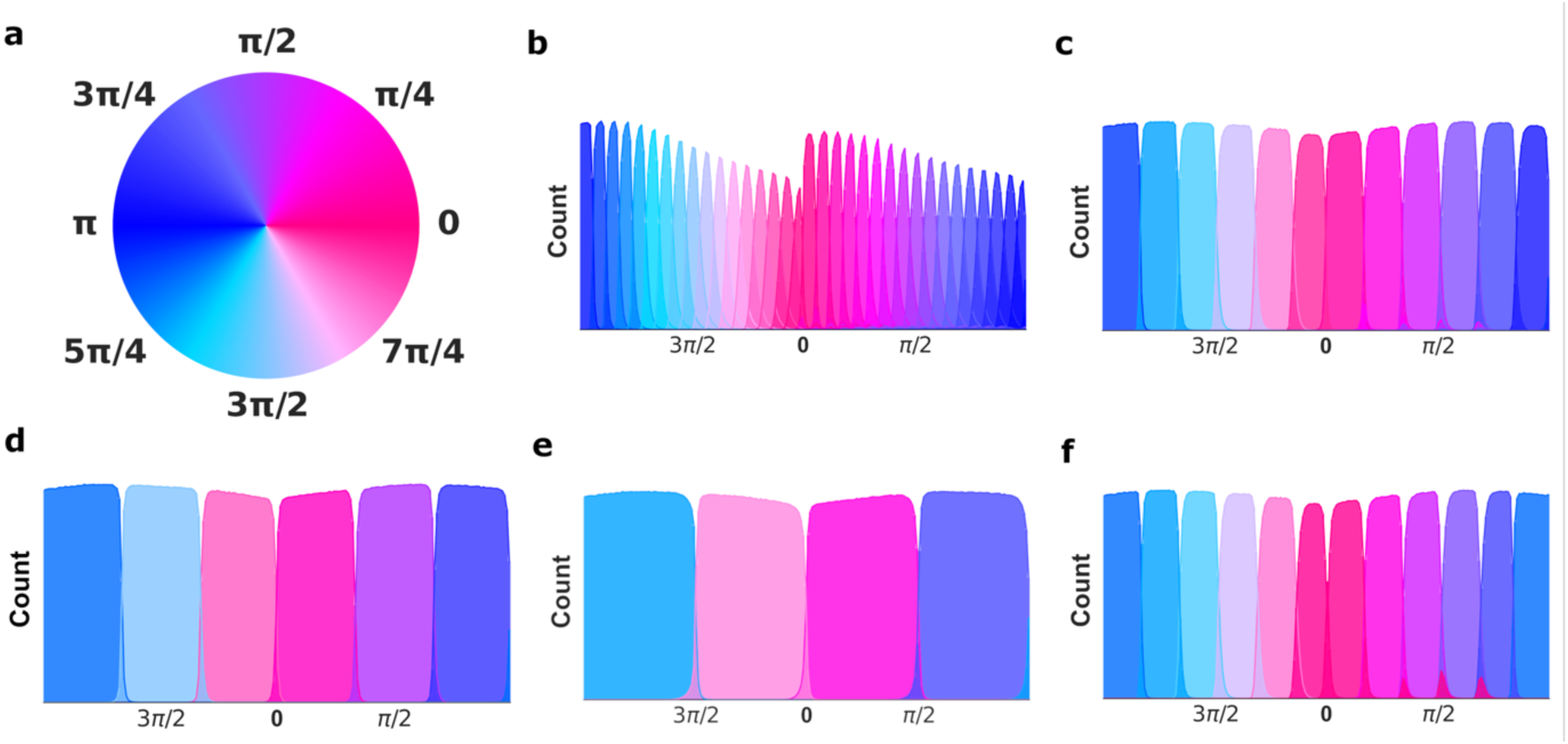
Distribution of instantaneous phase for parcels per community for the motor task fMRI data (mode 1). Data from all subjects, timepoints and parcels were aggregated. (a) Instantaneous phase values are represented by colours on the unit circle. (b) 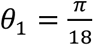, (c) 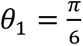, (d) 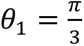, (e) 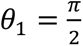, (f) 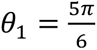. For all cases: *θ*_2_ = −*θ*_1_. Parcels are numbered as they appear from the left (−*π*) to right (*π*).

There is no known ground truth as to which degree of phase synchronization of BOLD-based time-series constitutes true cooperative processes between different brain regions. To show the applicability of the method to fMRI data, we made an arbitrary choice 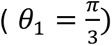. It should be noted that the general principles for the clustering algorithm as described above holds for other choices of thresholds as well. For convenience, communities were numbered from left to right as they appear in Figure 3. In both the lower and higher modes, the bulk of the parcels was divided into six communities. The six major communities had similar size measured as percentage of parcels across time and subjects (mode 1: mean 16.6% of parcels per community (total number of parcels = 436), SD = 0.6%, mode 2: mean 16.6%, SD = 0.4%,). However, a small number of parcels were assigned to one additional community (community number 7). The extra community represented a very small fraction of total amount of data. The number of parcels across time and subjects assigned to community 7^th^ community was 0.2% (mode 1) and 0.4% (mode 2). The occurrence of the 7^th^ community was correlated to the task paradigm block design in both modes (Suppl. Fig. S6). The relationship was particularly strong in mode 2 where it was less likely to appear during the cues than during task-blocks.

Figue 4 illustrates the average degree of separation between communities at the level of individual subjects and points in time. As shown in Figure 4, neighbouring communities have, on average, a segregation of cos(*x*) = 0.46, where x ≈ 63° and sin(*x*) = 0.86, were x ≈ 31°. (For standard deviations, see Suppl. Fig. S5).

**Figure 4.**
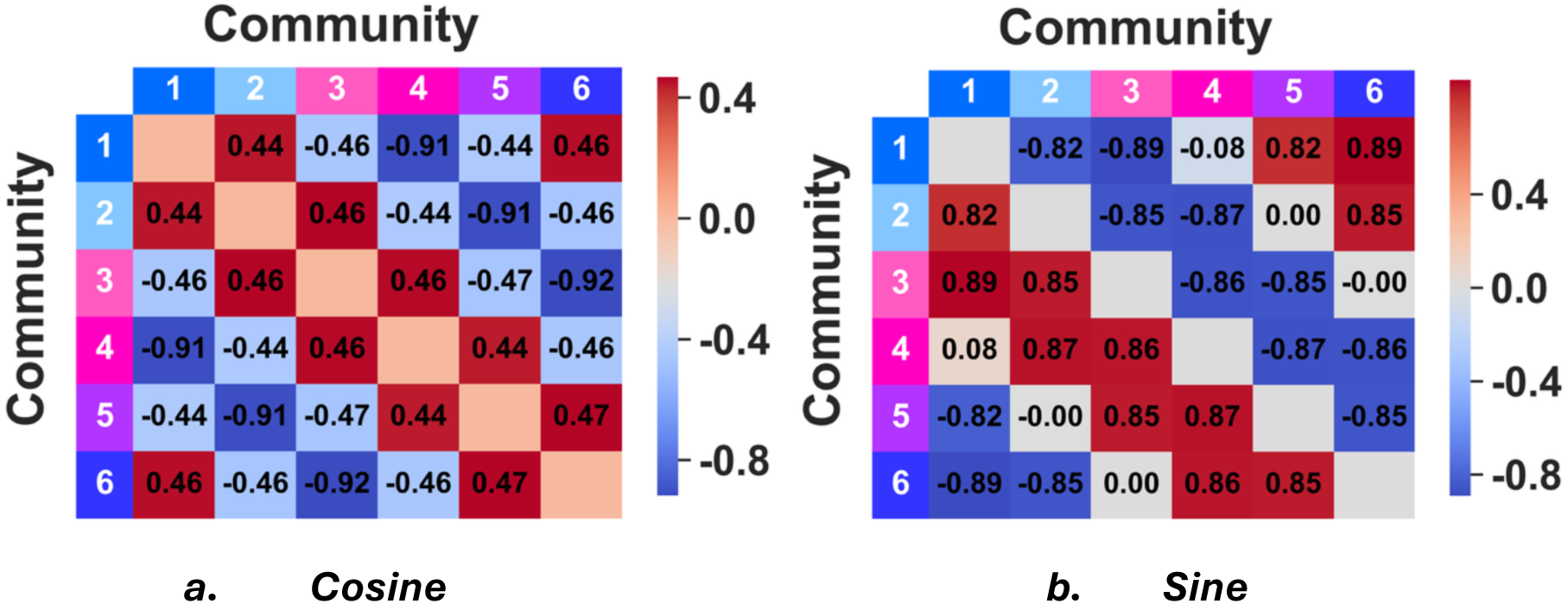
Illustration of community separation in terms of pairwise parcel phase differences (mode 1) between different communities (averaged across time points and subjects). (a) The mean cosine of the instantaneous phase difference (range [-1,1]) between pairwise parcels across communities (b) mean sine of the instantaneous phase difference (range [-1,1]) between pairwise parcels in the compared communities. The estimates of the standard deviations and corresponding results for mode 2 are provided in the Supplementary Material (Suppl. Fig. S7).

### 3.3 Network-specific cross-subject phase synchronization

To estimate the similarity in timing of the whole brain response to the motor task (both modes), we estimated the degree of cross-subject phase synchronization (see methods). In brief, it is defined as the percentage of subjects that are in synchrony with respect to community membership for a given point in time and parcel. The concept of cross-subject phase synchronization hinges on the fact that the algorithm assigns parcels to communities which span a limited phase range across time and subjects. Thus community labels are consistent (Fig. 3).

In the lower-frequency band (mode 1), Figures 5a, c and 6a show strong cross-subject phase synchronization during the five motor task blocks in the somato-motor network (SOM) parcels that are directly involved in the execution of the movements (left/right foot, left/right hand, and tongue movement). In the higher frequencies range (mode 2, Figs. 5b, d and 6b), cross-subject phase synchronization was observed most strongly and extensively in the visual (VIS) and attention networks (ventral, VAN, and dorsal DAN). Thus, our results for the higher frequency band (Fig. 6b) suggests a synchronization of the attentional networks that was independent of the specific motor movement task being executed. Rather, synchronization in the attentional networks seemed to reflect the specific timing and structure of the task block design. Moreover, higher frequency synchronization was also observed for parts of the DMN (default mode network), FPN (fronto-parietal network), SOM and TH (thalamic) networks, but to a considerably lesser extent. There is more uncertainty in the community assignment at certain sections of a timeseries (intersection between the community phase distributions Figs. 3 b-f), reflecting phases where transitions between communities are more common. This is seen as periods of lower synchronization interspersed within blocks of high synchronization creating a dotted appearance of peak synchronization in Figures 5a-b. In Figures 6a-b, cross-subject phase synchronization is depicted separately for each of the six main communities.

**Figure 5.**
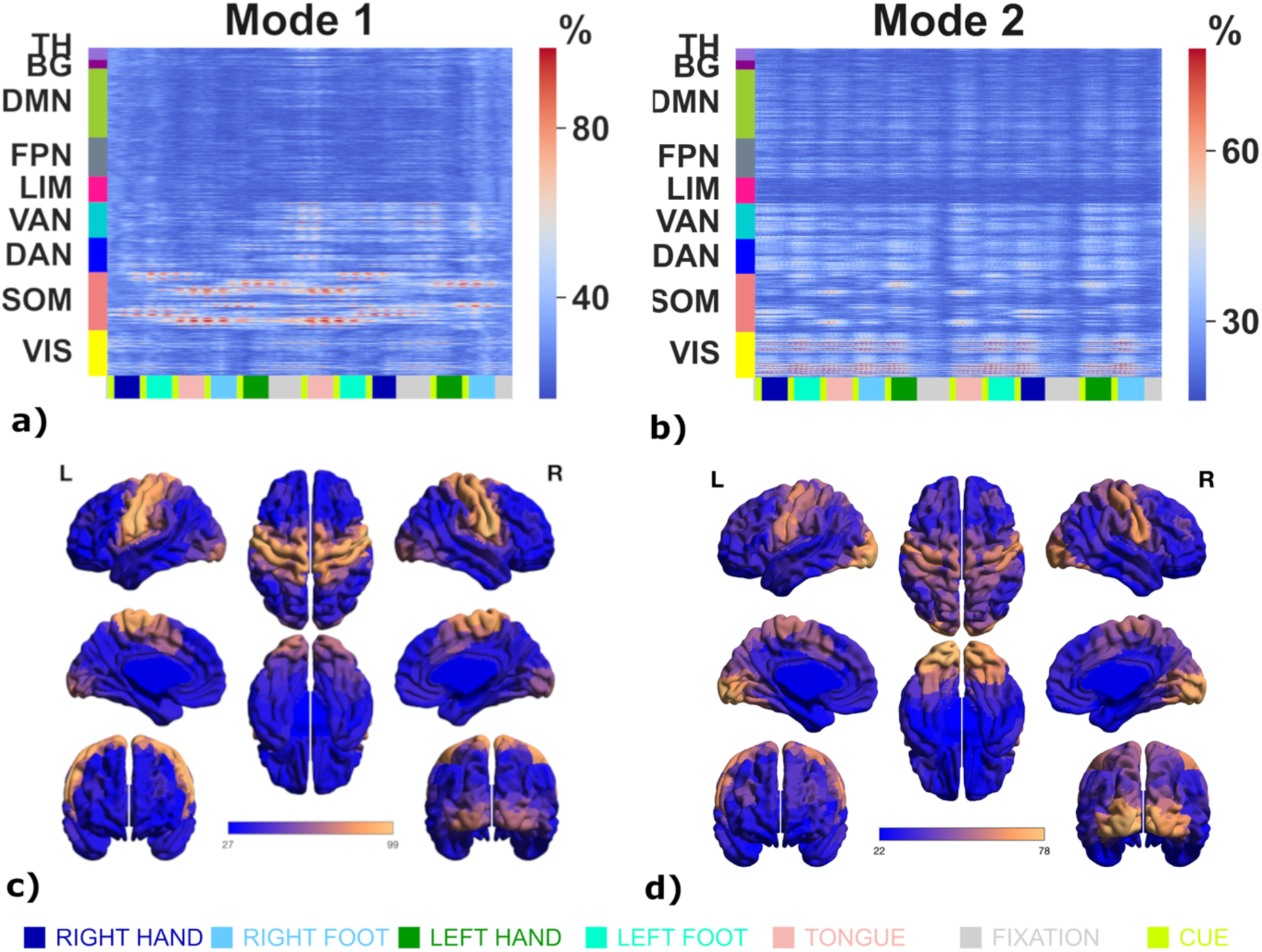
The maximum (computed across all 6 communities) of cross-subject phase synchronization (percentage of subjects) over time for (a) mode 1 (lower frequency range, max value 99%) and (b) mode 2 (higher frequency range, max value 77%) during the fMRI motor task. Panels (c) and (d) shows the maximal synchronization values projected onto the brain surface. An account of cross-subject synchronization in each community rather than the maximum values, is given in Figure 6.

**Figure 6a.**
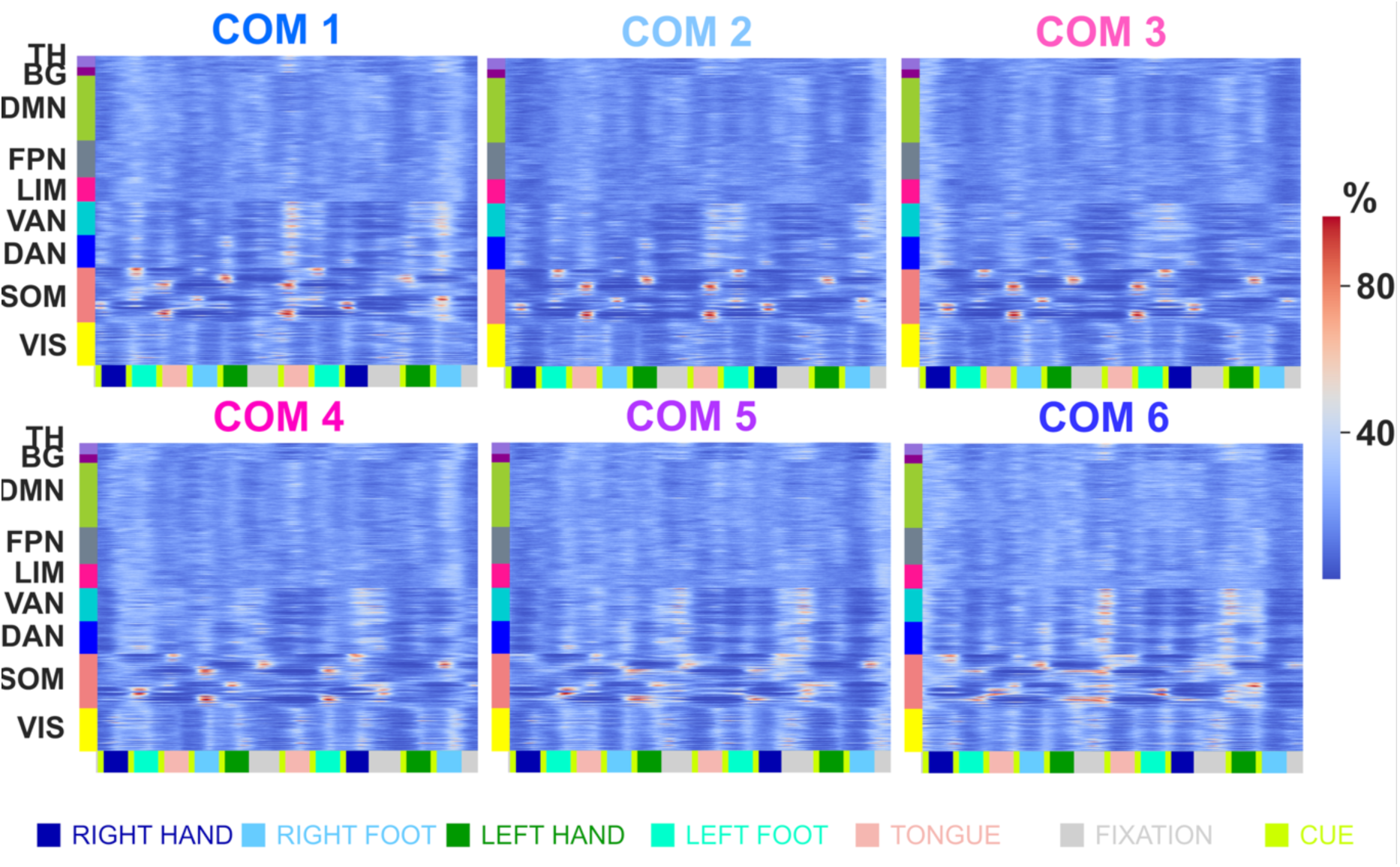
In this figure we show the degree of cross-subject phase synchronization for each community separately (COM1-COM6) for the lower frequency range (mode 1). Results pertaining to the higher frequency range (mode 2) is provided in Figure 6b. Colour coding reflects for each parcel the percentage of subjects residing in a particular community (COM1-COM6) at each timepoint, range [0,99 ]%. A maximum degree of synchronization (99% of all subjects) was observed in tongue motor area in the right hemisphere during the first tongue motor block (COM 3) but similar peaks of maximum cross-subject synchronization values were seen in all communities [86,99] % during tongue movement.

**Figure 6b.**
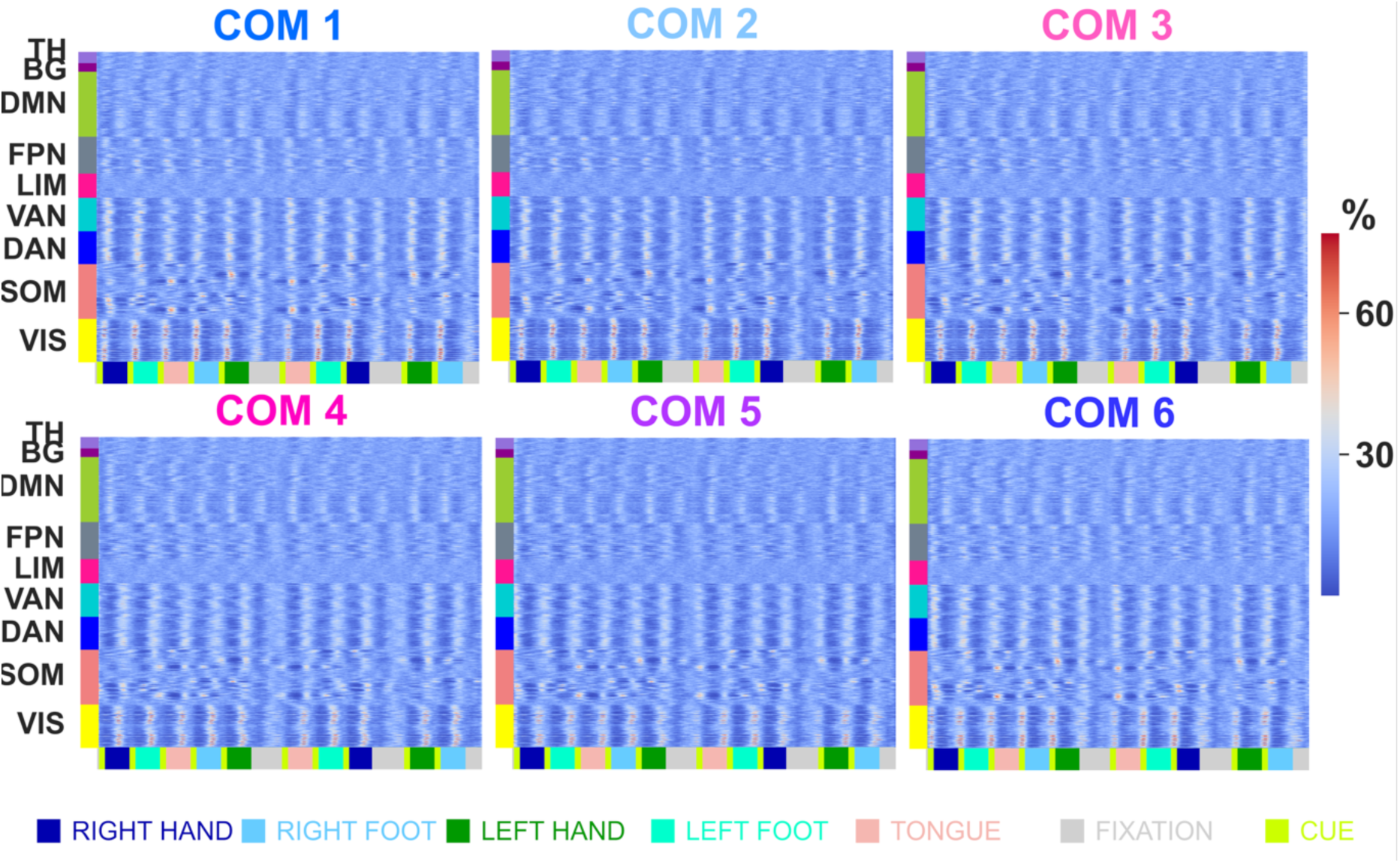
This figure complements the cross-subject phase synchronization results shown in Fig. 6a but shows the results for the higher frequency band (mode 2). Colour coding reflects for each parcel the percentage of subjects residing in a particular community (COM1-COM6) at each timepoint, range (0 - 77) %. The range of maximum cross subject phase synchronization was (67 – 77) %.

Figures 7-8 show how cross-subject phase synchronization timeseries relate in time to each other within single parcels. In Figure 7, the main motor parcels responsible for executing the motor tasks (mode 1) are used as an example. A parcel from the dorsal prefrontal cortex (PFC) is also included for comparison. Further examples, including a parcel located in the visual cortex (VIS), for mode 2 are shown in Figure 8.

**Figure. 7.**
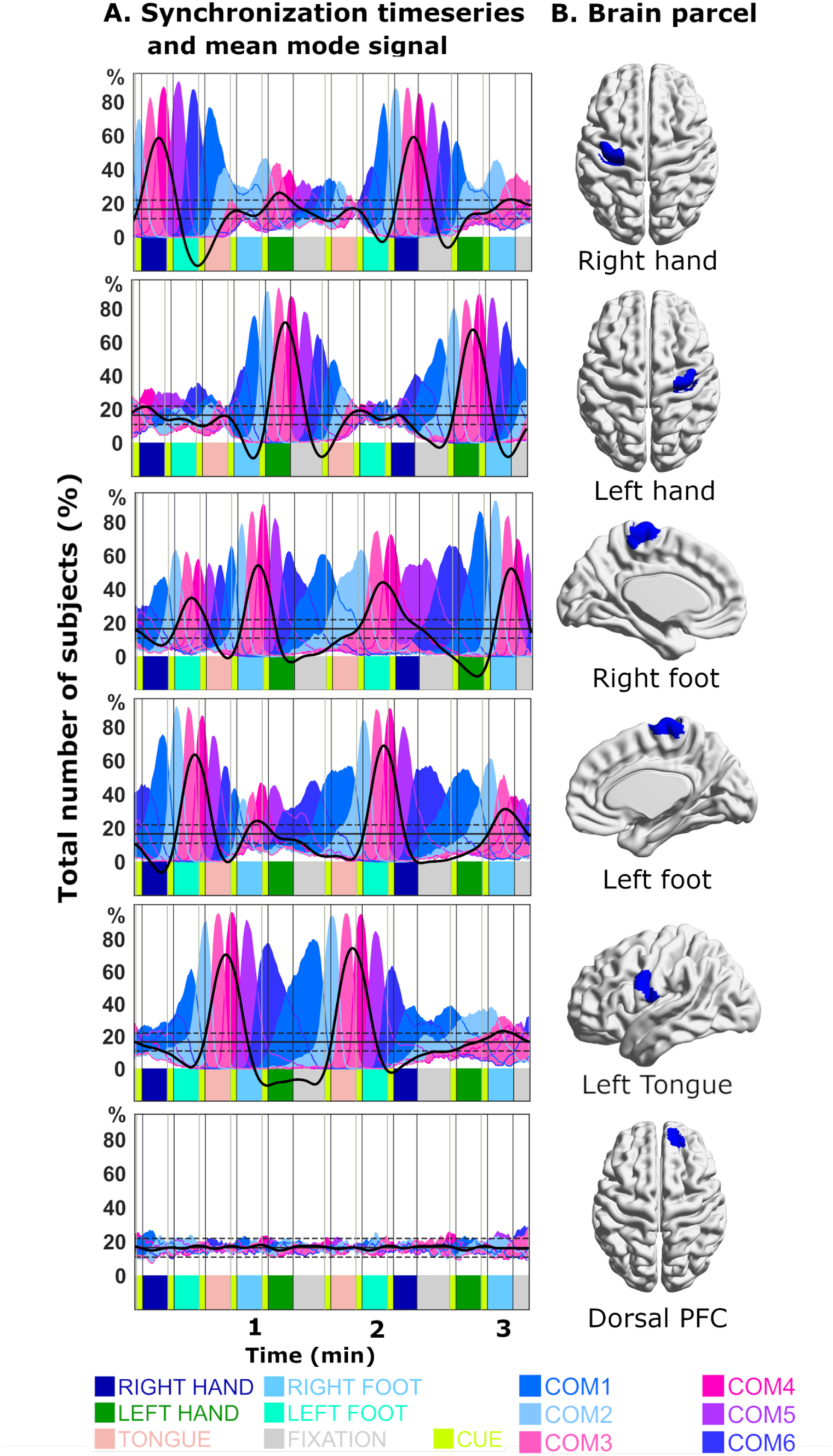
(a) Examples of degree of cross-subject phase synchronization as a function of time for five motor parcels involved in the motor task (Right tongue motor parcel not shown) for the lower frequency range (mode 1). A parcel in dorsal pre-frontal cortex (PFC) that did not show synchronization is also depicted for comparison. Black horizontal line corresponds to mean cross-subject phase synchronization for the aggregated phase scrambled data. Dotted lines correspond to ± 3*SD* in the distribution of phase scrambled data (See suppl. Fig. S3a). Mode 1 mean (across subjects) timeseries are overlayed in black. Mode timeseries have arbitrary units and are cantered at the mean for the phase scrambled cross-subject synchronization data. (b) Anatomical localisation of the parcels.

**Figure. 8.**
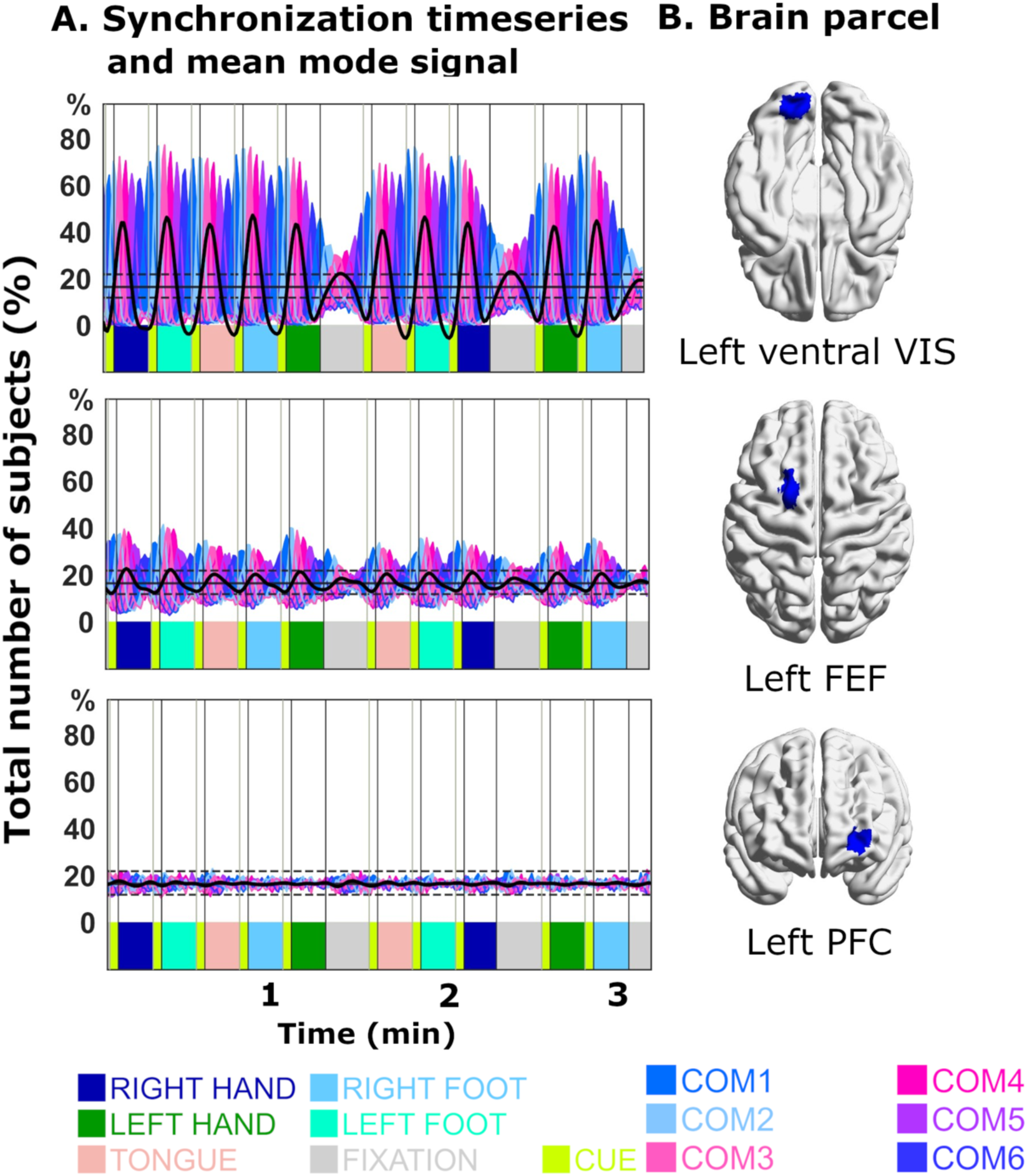
(a) Example of cross-subject phase synchronization timeseries in mode 2. Three parcels in three different networks are depicted: visual network (upper row), dorsal attention network (DAN, Left FEF4, middle row) and DMN (Left prefrontal cortex, PFC, lower row). Black horizontal lines correspond to mean cross-subject phase synchronization for the aggregated phase scrambled data in mode 2, i.e. 16.7 %. Dotted lines correspond to ± 3*SD* of the distribution of phase scrambled data (See suppl. Fig. S3b). Mode 2 (normalized) timeseries are overlayed in black. Mode timeseries have arbitrary units and are cantered at the mean for the phase scrambled cross-subject synchronization data. (b) Anatomical localisation of the parcels.

### 3.4 Integration and segregation

Since the relationships of parcels within and between communities are well defined by phase, the relative sizes of the communities can be used as measures of integration and segregation (see methods). Integration reflects positive phase relationships and segregation negative phase relationships. We analysed the whole-brain level and individual RSN:s to quantify the tension between integration and segregation throughout the motor task for the two modes. Statistical thresholds for significant results were calculated from the phase scrambled data (see methods).

At the whole-brain level, we observed distinct changes of community size as a function of time (Figure 9). For the lower frequency band (mode 1) shown in Figure 9a, we observed a “bimodal segregation” state (see methods, i.e. two communities that are maximally separated in phase are dominant) that lasts for about 20 seconds. This initial phase is followed by a “single domination” state (i.e. one community is substantially larger than the others, albeit with varying community sizes) that lasts throughout most of the remaining time of motor task experiment (Fig. 9a). Towards the end (last ∼20s), the “bimodal segregation” state reappears. For the higher frequency band (mode 2), the “single domination” state was observed throughout the motor task experiment with fluctuations in community sizes that are synchronized to the temporal design of the experiment (Fig. 9b).

**Figure 9.**
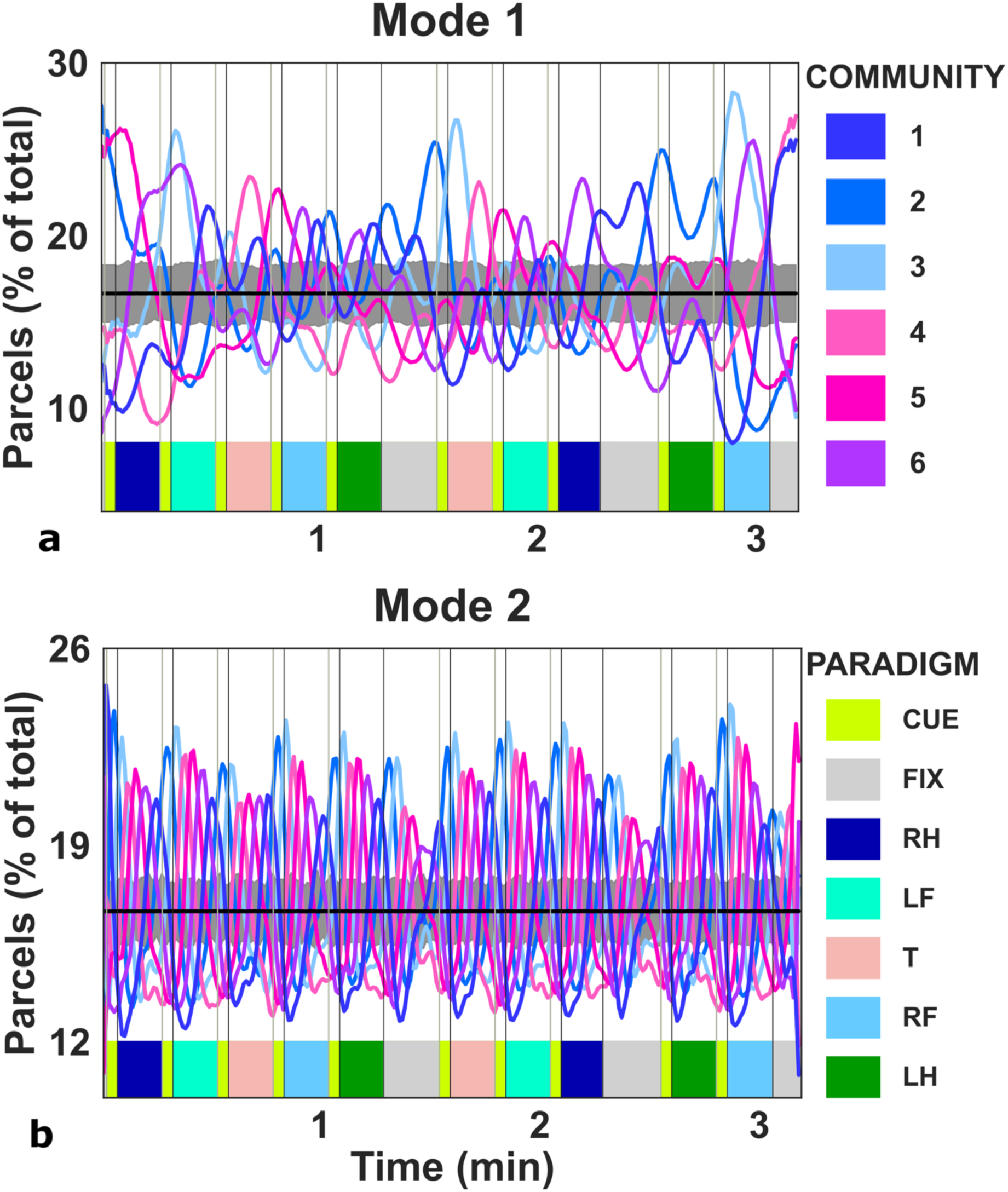
Average size of communities of parcels (percent of the total number of parcels) as a function of time (averaged across subjects). The black horizontal line represents the mean (across subjects) of the size of the six communities in the phase scrambled data. Gray shades represent ± 3*SD* of mean size fluctuations in the phase scrambled data. (a) mode 1, (b) mode 2 (FIX = fixation, RH = right hand, LF = left foot, T = tongue, RF = right foot, LG = left foot).

To get a clearer picture of what happens within the nine canonical resting-state networks (visual, somato-motor, dorsal and ventral attention, fronto-parietal, default mode, limbic, thalamic and basal ganglia networks) and how they contribute to whole-brain changes in integration/segregation, we computed the relative engagement of parcels (within each RSN) for each community at all points in time. The results are summarized in Figure 10 (lower frequency band, mode 1) and Figure 11 (higher frequency band, mode 2).

**Figure 10.**
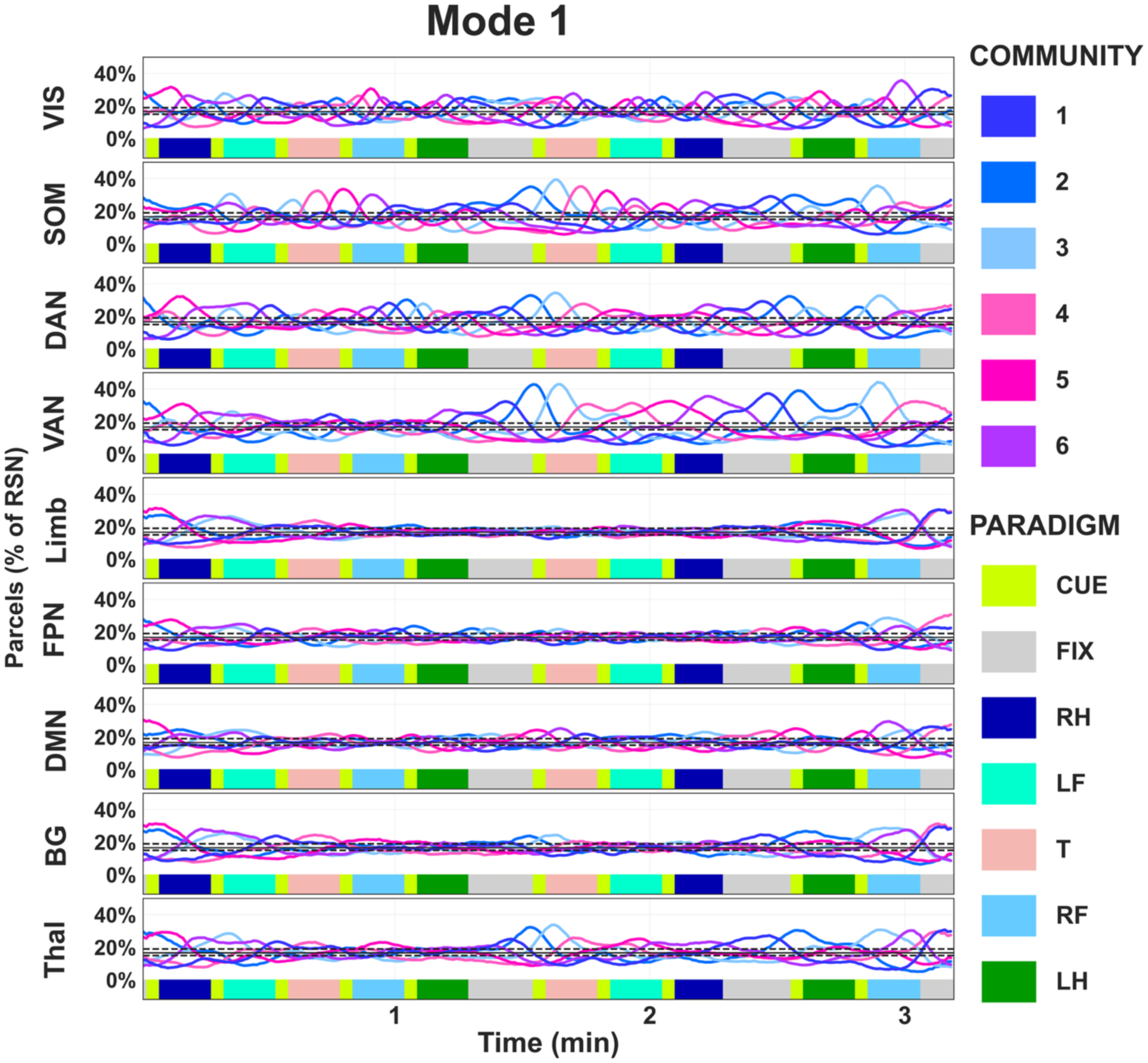
Relative contribution of parcels across time for the motor task with respect to the six major communities and divided based on the nine canonical RSNs (lower frequency range, mode 1). The y-axes represent the average (across subjects) percentage of the total number of parcels residing in each community for all canonical resting-state network in time. Statistical significance threshold +/-3 SD calculated from phase scrambled data as are depicted as dotted lines (FIX = fixation, RH = right hand, LF = left foot, T = tongue, RF = right foot, LG = left foot).

**Figure 11.**
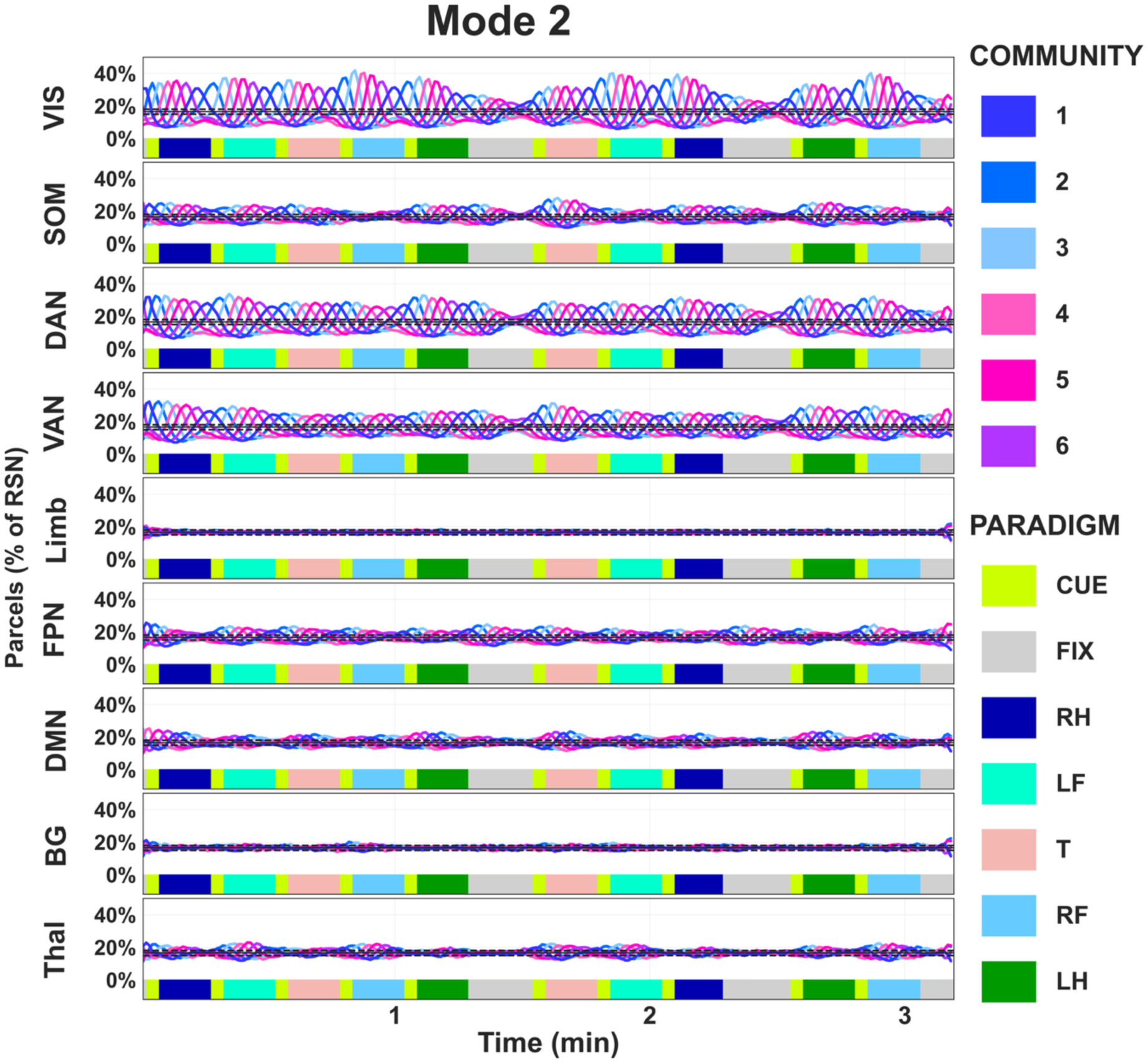
Relative contribution of parcels across time for the motor task with respect to the six major communities and divided based on the nine canonical RSNs (higher frequency range, mode 2). The y-axes represent the average (across subjects) percentage of the total number of parcels residing in each community for all canonical resting-state network in time. Statistical significance threshold +/-3 SD calculated from phase scrambled data as are depicted as dotted lines.

For the low frequency band (mode 1), a “bimodal segregation” community pattern was present at the beginning and end of the motor task run for the Limbic, Thalamic and BG networks (Fig. 10). The bimodal segregation behaviour was particularly prominent at the beginning and end of the motor task for the Limbic network. Regarding the time-period in-between the beginning and end, parcel community assignments for the canonical RSNs directly engaged by the task paradigm (VIS, SOM, DAN, VAN) was mostly of the “single dominating” type interspersed by short periods of “equalized” states. The RSNs that showed little engagement in the task paradigm (Limb, DMN, FPN, Thal, BG) where predominantly in a “equalized” state throughout. The “equalized state” corresponds to the non-synchronization of these networks. This means that community sizes equal out across subjects.

The results obtained for the higher frequency band (mode 2) shown in Figure 11 suggest that a single dominating community characterizes the parcel engagement in the visual, dorsal and ventral attention networks and to a lesser extent, the somato-motor network. The pattern of community switching time-locked to the task paradigm was the same. Thus, one can draw the conclusion that the patterns seen at the whole-brain level in mode 2 (Fig. 9b) to a large extent can be explained by the activity of visual and attention networks.

## 4. Discussion

We have developed an innovative approach that derives time-resolved communities, or networks, from parcellated fMRI data. These communities are delineated using integration and segregation thresholds, which partition parcels based on the instantaneous phase of the mode time series. The communities maintain consistency in phase internally and are distinctly separated in terms of the distributions of the instantaneous phase of parcels across time and subjects (Fig. 3-4). Since each community label reference a limited phase range, the method offers a means to assess the degree of synchronization of brain regions both intra- and inter-individually over time. Phase-scrambled data were employed to establish statistical thresholds of significance. Although we have selected an arbitrary cut-off for significance, most results substantially exceed this threshold.

Since signal phase is independent of amplitude, our method also captures synchronization where signal amplitudes are small, which are often ignored in conventional fMRI analysis methods. That being said, it is still debatable whether small amplitude fluctuations should be considered meaningful in fMRI data analyses (Cordes et al., 2001). Importantly, the method does not only capture phase synchronization during peak motor activations but also through cycles of positive and negative amplitude shifts (Suppl. Fig. S8). Since the method respects the temporal order of the timeseries, it can be used for a fine-grained analysis of differences in timing between brain regions (Bostami et al., 2025; Cole et al., 2016; Mitra et al., 2014; Reid et al., 2019).

An important feature of the proposed method is the ability of the VMD algorithm to decompose the band-pass filtered BOLD signal into meaningful modes, i.e. frequency ranges, that reflect distinct aspects of the data. For the motor task dataset used here, the VMD decomposition of the data entailed that phase synchronization, expressed as community membership, for the attention and visual networks resided in the higher frequency range (mode 2, see Figures 5, 6, 8 and 11). In contrast, changes in community membership related to specific changes in the somato-motor network was encapsulated by the slower frequency range (mode 1, see Figures 5, 6, 7 and 10).

The results shown here pertaining the two different modes can be related to previous research examining the properties of brain activation/connectivity as a function of frequency of the BOLD-signal. Previously, the low-frequency fluctuations of the BOLD signal have been divided into 4 bands: slow-5 (0.01–0.027 Hz), slow-4 (0.027–0.073 Hz), slow-3 (0.073–0.198 Hz), slow-2 (0.198–0.25 Hz) and slow-1 (0.5 – 0.75 Hz) based on the subdivisions of the full spectrum of brain oscillations from neurophysiological recordings with EEG (Buzsáki & Draguhn, 2004; Penttonen, 2003) (Di Martino et al., 2008; Zuo et al., 2010). With a sampling rate of 0.5 Hz, BOLD-fluctuations in the grey matter have shown the strongest fluctuations in the slow-4 and slow-5 frequency bands (Zuo et al., 2010). Using even higher sampling rates (TRs), it has been shown that RSN connectivity also resides in the higher frequency bands (Boubela et al., 2013; Gohel & Biswal, 2015; Lee et al., 2013; Niazy et al., 2011).

The frequency range designated as mode 1 in our study approximately corresponds to the slow-5 band while mode 2 can be seen as an aggregation of the slow-4 band and parts of the slow-3 band. In electrophysiological recordings, slower frequencies recruit larger networks and higher frequencies are limited to smaller localized networks (Buzsáki & Draguhn, 2004; Varela et al., 2001). Previous work in fMRI resting state data using sampling rates similar to or higher than that of the HCP-data (1.4 Hz) used in this study, have shown that RSN connectivity is present in all bands with similar spatial extent but with band specific connectivity (Gohel & Biswal, 2015; Salvador et al., 2008; Thompson & Fransson, 2015). Our results of a mode specific segregation of network activity directly relates to the task block design and the execution of the simple motor tasks. A different type of paradigm that is less predictable will likely not induce the same clear separation between attention networks and task specific network activation into different modes. We would also not expect the same segregation in resting state data. More likely, in line with previous work, activity of all networks will be represented approximately equally in all modes for resting state fMRI data.

### 4.1 Integration and segregation

The concepts of integration and segregation in fMRI studies commonly refer to two types of (whole) brain states (Bassett et al., 2011b, 2013; Betzel et al., 2016, 2016; Deco et al., 2015; Fukuda et al., 2016; Fukushima, Betzel, He, De Reus, et al., 2018; Fukushima, Betzel, He, Van Den Heuvel, et al., 2018; Handwerker et al., 2012; Shine et al., 2016; Sporns, 2013; Thompson & Fransson, 2015; Zalesky et al., 2014). Integrated states are periods characterized by relatively high connectivity between networks. Conversely, segregation is defined as weak connectivity between networks and high connectivity within networks (together referred to as modularity). The intrinsic activity of the brain spontaneously fluctuates between periods of integration and segregation (Zalesky et al., 2014). Task demands tend to induce integration (Shine et al., 2016). In the present (and previous (Fransson & Strindberg, 2023; Strindberg et al., 2021)) work, we use integration and segregation to refer to relationships between parcels (within communities) or communities defined by phase. Integration is present on a gradient from strong positive in-phase relationships (based on the chosen threshold) within communities, to weaker positive in-phase relationships between communities. In this context, segregation refers to a negative phase relationship between communities that also exists on the opposite gradient (from strong negative to weak negative phase relationships). Within this framework, integration and segregation are simultaneous in the brain and describe the relationships both within and between communities in a graded fashion, but their relative strengths may vary across time. Thus, our reasoning and method does not contradict to the more common use of the concepts but rather endorses a somewhat more nuanced view.

We analysed the level of time-resolved integration and segregation by means of considering the average sizes (in terms of parcels) of communities (see Fig. 9). At a whole-brain level for the lower frequency band (mode 1), the start and end periods of the motor task paradigm were associated with a split of at least half of the parcels into two maximally segregated communities (on average). In between, for most of the time, one community was dominating in size, albeit the extent of dominance was variable (Fig. 9a). One could speculate that the “bimodal segregation” of community assignment observed at the beginning and towards the end of the motor task paradigm is related to anticipatory processes. However, to confirm or refute this hypothesis necessitates further work. At the level of the canonical RSNs, the bimodal segregation state seen on the whole-brain level at the beginning and end of the motor task paradigm in the lower frequency band (mode 1) (Fig. 9a), was also present within the subcortical networks (Thal, BG) and the Limbic network and to a lesser extent the other networks (Fig. 10).

In the higher frequency band (mode 2), a single community dominated throughout the paradigm, but less pronounced during fixation blocks (Fig. 9b). The pattern was highly repetitive. Independent of the specific task (except for the fixation blocks), neighbouring communities (see also Figs. 3a and d) followed one another counterclockwise as they appear on the unit circle in sync with the block-design (Figs. 8 and 9). The strongly repeating pattern of a single dominating community in mode 2 was largely due to the coordinated and continuous recruitment of the visual (VIS) and the dorsal (DAN) and ventral (VAN) attention networks throughout the task block-design (Fig. 11). These canonical networks remain engaged and synchronized across subjects throughout the cycles of positive and negative amplitude shifts (see also Figures 5b, 6b). This finding highlights the importance of considering both positive as well as negative activations (i.e. deactivations), and the gradual shift between them, rather than only the peak positive BOLD amplitudes in fMRI studies.

### 4.2 Limitations and properties of the VMD algorithm

Our method relies on the mode timeseries extracted from bandpass filtered data using the VMD algorithm. How well these modes together reflect the unfiltered timeseries enforces limits of reliability of our results and the conclusions one can draw from them. Moreover, our results may also be influenced by our choice of bandpass filter parameters and the number of a priori defined modes. However, the effect of the band-pass filter was small (Suppl. Fig. S4). Of note, the VMD algorithm uses the Fourier transform. If the VMD algorithm is forced to decompose more signal modes than what are potentially present in the data, it can create false positives which presents themselves as higher frequency harmonics in the decomposed signal modes. This fact must be contrasted with the brain response elicited by a highly regular task-design such as the block-design. In the motor-task data used here as example data, the power spectra of the two modes closely resemble the power spectrum of the band-pass filtered and raw data (Suppl. Fig. S4). Harmonics of the first (and largest) peak (P1) at 0.016 Hz (Suppl. Fig. S10) were seen at 0.032 Hz (2*P1, peak 2), 0.047Hz (3*P1, peak 3). P4 which was the largest peak in mode 2, was close to being a harmonic too (i.e. a “perfect” harmonic counting three decimal places would have been 0.063 but P4 was 0.068, see Suppl. Fig S10). Given that P4 clearly was evoked by the block design (3+12 seconds for cue+ task blocks and 15s for fixation blocks) it unlikely to be a harmonic of the base peak. These peaks were also seen in the raw data (Suppl. Fig. S4, S10). Thus, they were not artifacts from the VMD-algorithm but features of the data. With regard to mode 1 (lower frequency band), the relationship between the peaks in the power spectrums and the task-evoked activity is not as clear cut; The period of highest synchronization across subjects of the task specific parcels in mode 1 was ∼45*s* (Fig. 5a) which lies in between P1 and P2 in the power spectrum (Suppl. Fig. S10). The degree of cross-subject phase synchronization is governed by the individual modal signal time courses. The two peaks in mode 1 (Suppl. Fig. S10) seem to reflect the intrinsic activity of the brain where task specific activity is present but not the only, or even most important, contribution to the corresponding power spectrum (Suppl. Fig. S5a). This is in line with what is already well known about the brain, i.e. task-evoked activity represent only a fraction of the energy consumed by the intrinsic activity of the brain (Raichle et al., 2001) and that evoked activity modulates intrinsic activity (Cole et al., 2014, 2016).

Higher order harmonics of signals of interest are more commonly discussed in the M/EEG literature. They are by some authors proposed to be a sign of entrainment in systems of non-linear dynamics (Norcia et al., 2015). Entrainment refers to the idea that intrinsic rhythms are recruited to perform the task at hand (Keitel et al., 2014, 2022). This is contrasted with the idea that evoked activity “provokes” the generation of the corresponding rhythms. Whether entrainment is a real phenomenon is highly debated and methodologically difficult to examine (Duecker et al., 2024; Haegens, 2020; Meyer et al., 2020). The presence of harmonics in this dataset was just an observation. The paradigm per se was not designed to investigate the possibility of entrainment in BOLD signals or other intrinsic mechanisms that would explain this phenomenon. Nor have we used correction for physiological noise (heart rate and respiration) to rule out aliasing of heart rhythms and their potential harmonics (Cordes et al., n.d.; Glover et al., 2000). However, the apparent presence of higher order harmonics in BOLD motor task data raises some interesting questions that should be addressed by further research.

While we tried to reduce the degree of mode mixing in the VMD-step by conducting a pre-VMD band-pass filtering of the data, the choice of the bandpass filter parameters might to some extent influence the results. However, given that the peaks in the power-spectra remained intact in relation to the raw data (Supp. Fig. S4), this effect is likely minor. Our choice of decomposing the bandpass-filtered BOLD signal into two variational modes by the VMD algorithm was guided by previous work using the (CE)EMD algorithms which estimates the number of modes directly from the data. An in-depth analysis of the most appropriate number of modes for a particular data must be guided by the sampling rate and potentially task design. We do not claim to have found the optimal combination of band-pass filter and number of VMD modes. These issues warrant further research. While the higher frequencies in the BOLD-signal have low power due to the inherent low-pass filter created by the hemodynamic response function, in future work it would be interesting to investigate the effects of using a wider band-pass filter and allowing for a higher number of VMD modes applied to more complex task paradigms as well as for resting-state data.

### 4.3 Limitations of the phase clustering algorithm

An obvious weakness of our method is the elevated uncertainty of community assignment at phase values close to the intersections between neighbouring community distributions (Fig. 3). This limitation is perhaps most clearly visible in Figures 5 a, b where intermittent peaks of maximum cross-subject synchronization are interspersed with short periods of relatively less synchronization. Depending on the research question at hand, this might or might not be a problem. If it is important to define the maximum cross-subject phase synchronization at all time-points, the problem can be mitigated by calculating additional sine-based communities. Using the sine part of the instantaneous phase signal instead of cosine as a basis in the phase cluster algorithm will create communities where the intersection between the community distributions is shifted by 90 degrees. These two sets of communities, with cosine and sine as a basis for clustering respectively, will together yield a more complete representation of maximum cross-subject phase synchronization. This means that if the maximum synchronization was calculated based on both sets of communities, the periods of high synchronization in Fig. 5 would have a more continuous appearance. The reason is that the phase range of elevated uncertainty are non-overlapping.

The algorithm optimizes for integration (i.e. phase consistent communities, *thr1*) and maximal segregation (if present, *thr2*). The choice of integration/segregation thresholds (*thr1* and *thr2*) directly controls the number of communities that are formed (Fig. 3). Thresholding per se introduces an artificial boundary that might not (and is most cases is not) biologically meaningful. However, it is helpful in sorting and quantifying complex information.

In the results presented here, the average cosine of phase difference (cos (*θ*)) between neighbouring communities was only slightly lower than the chosen *thr1* (0.46 vs 0.5) (Fig. S4). These results show the tight relationship between neighbouring communities, and it highlights the fact that there is rarely an optimal way to find community structures for the highly interconnected networks existing in the brain. Whether the interconnectedness between communities is an obstacle or an opportunity really depends on the research question at hand. For example, if a tracking of the gradual changes in network configurations along the gradient between being integrated versus segregated is needed, the close but well-defined relationships between neighbouring communities as shown here is useful.

The algorithm is biased in the sense that it always finds the parcel with the most negative phase value first in each iteration. This is by design to create reproducible communities. If one wants to optimize for the largest community primarily, this might not be the ideal choice. We did explore a version of the algorithm that optimizes for the largest community as the first step. This outcome was however obtained at the cost of reproducible communities and hence synchronization across time and subjects. While the clustering algorithm presented here sets constraints on the consistency of phase within the community and in relation to a maximally segregated community (if present), it has size as the secondary optimization factor (see Suppl. Fig. S9 for distributions of community sizes for the two modes). The permutation option of the algorithm optimizes this step further (not used in obtaining the results presented here).

### 4.4 Different uses of the phase clustering algorithm

For clarity, in our presentation of the phase clustering algorithm we have focused on one arbitrary choice of thresholds (thr1=cos(*θ*_1_), *thr*2 = −*thr*1, *w*ℎ*ere* 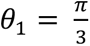) to illustrate the use of the algorithm to answer questions about synchronized responses to task paradigms. However, the algorithm can be used with different choices of thresholds. For example, the segregation threshold (*thr2*) can be omitted completely or set to a different value, i.e. (*θ*_2_ ≠ −*θ*_1_). It is the research question that must guide the choice of thresholds. The phase clustering algorithm is designed to find common structures in multiple sets of time-resolved data, i.e. where each set represent data sampled at different timepoints. However, an alternative would be to use it to find patterns of interest across multiple datasets for a single time-point only. While the algorithm is designed for data where approximately the full range of phase values are represented in each timepoint and each data set, it could be adjusted to fit data that don’t fulfil this set of criteria.

Although it is not explicitly demonstrated here, the method’s granularity allows for the estimation of both directionality and the strength of activity flow (Mitra et al., 2014; Reid et al., 2019). Directionality between individual parcels or networks activity could be assessed by calculating the timing of the peaks of cross-subject synchronization in the same community. The strength of the relationship could be estimated by assessing the level of cross subject synchronization. A strong “excitatory” relationship would be present during time-periods when synchronization was high in both the leading and lagging peaks. In the same way one could assess a potentially “inhibitory” flow by assessing the timing of the peaks of synchronization in maximally segregated communities. The choice of thresholds and the granularity of the parcellation will influence the results as will the quality of the data.

In summary, we have presented a novel method which reliably identifies synchronization across subjects (if present) through periods of activation and deactivation. In the fMRI data set analysed here, cross-subject synchronization at times reached 99% of all subjects. The result from our study provides a complementary view on brain network integration and segregation and allows for the analysis of the temporal order of network recruitment within and between individuals.

## Code availability

The code described in this paper is available for use at https://github.com/TemporalBrain/PhaseClustering

## Author contributions

MS: Conceptualization, Methodology, Writing-original draft, Writing-reviewing and editing. PF: Supervision, Validation, Writing-reviewing and editing.

## References

Allen, E. A., Damaraju, E., Plis, S. M., Erhardt, E. B., Eichele, T., & Calhoun, V. D. (2014). Tracking Whole-Brain Connectivity Dynamics in the Resting State. Cerebral Cortex, 24(3), 663–676. 10.1093/cercor/bhs352

Andrews-Hanna, J. R., Reidler, J. S., Sepulcre, J., Poulin, R., & Buckner, R. L. (2010). Functional-Anatomic Fractionation of the Brain’s Default Network. Neuron, 65(4), 550–562. 10.1016/j.neuron.2010.02.005

Bandettini, P. A., Wong, E. C., Hinks, R. S., Tikofsky, R. S., & Hyde, J. S. (1992). Time course EPI of human brain function during task activation. Magnetic Resonance in Medicine, 25(2), 390–397. 10.1002/mrm.1910250220

Barch, D. M., Burgess, G. C., Harms, M. P., Petersen, S. E., Schlaggar, B. L., Corbetta, M., Glasser, M. F., Curtiss, S., Dixit, S., Feldt, C., Nolan, D., Bryant, E., Hartley, T., Footer, O., Bjork, J. M., Poldrack, R., Smith, S., Johansen-Berg, H., Snyder, A. Z., & Van Essen, D. C. (2013). Function in the human connectome: Task-fMRI and individual differences in behavior. NeuroImage, 80, 169–189. 10.1016/j.neuroimage.2013.05.033

Bassett, D. S., Porter, M. A., Wymbs, N. F., Grafton, S. T., Carlson, J. M., & Mucha, P. J. (2013). Robust detection of dynamic community structure in networks. Chaos: An Interdisciplinary Journal of Nonlinear Science, 23(1), 013142. 10.1063/1.4790830

Bassett, D. S., Wymbs, N. F., Porter, M. A., Mucha, P. J., Carlson, J. M., & Grafton, S. T. (2011a). Dynamic reconfiguration of human brain networks during learning. Proceedings of the National Academy of Sciences, 108(18), 7641–7646. 10.1073/pnas.1018985108

Bassett, D. S., Wymbs, N. F., Porter, M. A., Mucha, P. J., Carlson, J. M., & Grafton, S. T. (2011b). Dynamic reconfiguration of human brain networks during learning. Proceedings of the National Academy of Sciences, 108(18), 7641–7646. 10.1073/pnas.1018985108

Beckmann, C. F., DeLuca, M., Devlin, J. T., & Smith, S. M. (2005). Investigations into resting-state connectivity using independent component analysis. Philosophical Transactions of the Royal Society B: Biological Sciences, 360(1457), 1001–1013. 10.1098/rstb.2005.1634

Berger, H. (1929). Über das Elektrenkephalogramm des Menschen. Archiv für Psychiatrie und Nervenkrankheiten, 87(1), 527–570. 10.1007/BF01797193

Betzel, R. F., Fukushima, M., He, Y., Zuo, X.-N., & Sporns, O. (2016). Dynamic fluctuations coincide with periods of high and low modularity in resting-state functional brain networks. NeuroImage, 127, 287–297. 10.1016/j.neuroimage.2015.12.001

Billings, J. C. W., Medda, A., Shakil, S., Shen, X., Kashyap, A., Chen, S., Abbas, A., Zhang, X., Nezafati, M., Pan, W.-J., Berman, G. J., & Keilholz, S. D. (2017). Instantaneous brain dynamics mapped to a continuous state space. NeuroImage, 162, 344–352. 10.1016/j.neuroimage.2017.08.042

Biswal, B., Zerrin Yetkin, F., Haughton, V. M., & Hyde, J. S. (1995). Functional connectivity in the motor cortex of resting human brain using echo-planar mri. Magnetic Resonance in Medicine, 34(4), 537–541. 10.1002/mrm.1910340409

Bostami, B., Lewis, N., Agcaoglu, O., Turner, J. A., van Erp, T., Ford, J. M., Fouladivanda, M., Calhoun, V., & Iraji, A. (2025). Time-Varying Spatial Propagation of Brain Networks in fMRI Data. Human Brain Mapping, 46(2), e70131. 10.1002/hbm.70131

Boubela, R. N., Kalcher, K., Huf, W., Kronnerwetter, C., Filzmoser, P., & Moser, E. (2013). Beyond Noise: Using Temporal ICA to Extract Meaningful Information from High-Frequency fMRI Signal Fluctuations during Rest. Frontiers in Human Neuroscience, 7. 10.3389/fnhum.2013.00168

Breakspear, M. (2017). Dynamic models of large-scale brain activity. Nature Neuroscience, 20(3), 340–352. 10.1038/nn.4497

Bullmore, E., & Sporns, O. (2009). Complex brain networks: Graph theoretical analysis of structural and functional systems. Nature Reviews Neuroscience, 10(3), 186–198. 10.1038/nrn2575

Buzsáki, G., & Draguhn, A. (2004). Neuronal Oscillations in Cortical Networks. Science, 304(5679), 1926–1929. 10.1126/science.1099745

Cabral, J., Vidaurre, D., Marques, P., Magalhães, R., Silva Moreira, P., Miguel Soares, J., Deco, G., Sousa, N., & Kringelbach, M. L. (2017). Cognitive performance in healthy older adults relates to spontaneous switching between states of functional connectivity during rest. Scientific Reports, 7(1), 5135. 10.1038/s41598-017-05425-7

Calhoun, V. D., Miller, R., Pearlson, G., & Adalı, T. (2014). The Chronnectome: Time-Varying Connectivity Networks as the Next Frontier in fMRI Data Discovery. Neuron, 84(2), 262–274. 10.1016/j.neuron.2014.10.015

Chang, C., & Glover, G. H. (2010). Time–frequency dynamics of resting-state brain connectivity measured with fMRI. NeuroImage, 50(1), 81–98. 10.1016/j.neuroimage.2009.12.011

Cole, M. W., Bassett, D. S., Power, J. D., Braver, T. S., & Petersen, S. E. (2014). Intrinsic and Task-Evoked Network Architectures of the Human Brain. Neuron, 83(1), 238–251. 10.1016/j.neuron.2014.05.014

Cole, M. W., Ito, T., Bassett, D. S., & Schultz, D. H. (2016). Activity flow over resting-state networks shapes cognitive task activations. Nature Neuroscience, 19(12), 1718–1726. 10.1038/nn.4406

Cordes, D., Haughton, V. M., Arfanakis, K., Carew, J. D., Turski, P. A., Moritz, C. H., & Quigley, M. A. (n.d.). Frequencies Contributing to Functional Connectivity in the Cerebral Cortex in ‘“Resting-state”’ Data.

Cordes, D., Haughton, V. M., Arfanakis, K., Carew, J. D., Turski, P. A., Moritz, C. H., Quigley, M. A., & Meyerand, M. E. (2001). Frequencies contributing to functional connectivity in the cerebral cortex in ‘resting-state’ data. AJNR. American Journal of Neuroradiology, 22(7), 1326–1333.

Damoiseaux, J. S., Rombouts, S. a. R. B., Barkhof, F., Scheltens, P., Stam, C. J., Smith, S. M., & Beckmann, C. F. (2006). Consistent resting-state networks across healthy subjects. Proceedings of the National Academy of Sciences of the United States of America, 103(37), 13848–13853. 10.1073/pnas.0601417103

Deco, G., Ponce-Alvarez, A., Mantini, D., Romani, G. L., Hagmann, P., & Corbetta, M. (2013). Resting-State Functional Connectivity Emerges from Structurally and Dynamically Shaped Slow Linear Fluctuations. Journal of Neuroscience, 33(27), 11239–11252. 10.1523/JNEUROSCI.1091-13.2013

Deco, G., Tononi, G., Boly, M., & Kringelbach, M. L. (2015). Rethinking segregation and integration: Contributions of whole-brain modelling. Nature Reviews Neuroscience, 16(7), 430–439. 10.1038/nrn3963

Di Martino, A., Ghaffari, M., Curchack, J., Reiss, P., Hyde, C., Vannucci, M., Petkova, E., Klein, D. F., & Castellanos, F. X. (2008). Decomposing Intra-Subject Variability in Children with Attention-Deficit/Hyperactivity Disorder. *Biological Psychiatry*, Endophenotypes for Autism Spectrum Disorder and Attention-Deficit/Hyperactivity Disorder, 64(7), 607–614. 10.1016/j.biopsych.2008.03.008

Dragomiretskiy, K., & Zosso, D. (2014). Variational Mode Decomposition. IEEE Transactions on Signal Processing, 62(3), 531–544. 10.1109/TSP.2013.2288675

Duecker, K., Doelling, K. B., Breska, A., Coffey, E. B. J., Sivarao, D. V., & Zoefel, B. (2024). Challenges and Approaches in the Study of Neural Entrainment. The Journal of Neuroscience, 44(40), e1234242024. 10.1523/JNEUROSCI.1234-24.2024

Fan, L., Li, H., Zhuo, J., Zhang, Y., Wang, J., Chen, L., Yang, Z., Chu, C., Xie, S., Laird, A. R., Fox, P. T., Eickhoff, S. B., Yu, C., & Jiang, T. (2016). The Human Brainnetome Atlas: A New Brain Atlas Based on Connectional Architecture. Cerebral Cortex, 26(8), 3508–3526. 10.1093/cercor/bhw157

Fox, M. D., Snyder, A. Z., Vincent, J. L., Corbetta, M., Van Essen, D. C., & Raichle, M. E. (2005). The human brain is intrinsically organized into dynamic, anticorrelated functional networks. Proceedings of the National Academy of Sciences, 102(27), 9673–9678. 10.1073/pnas.0504136102

Fransson, P. (2005). Spontaneous low-frequency BOLD signal fluctuations: An fMRI investigation of the resting-state default mode of brain function hypothesis. Human Brain Mapping, 26(1), 15–29. 10.1002/hbm.20113

Fransson, P., & Strindberg, M. (2023). Brain network integration, segregation and quasi-periodic activation and deactivation during tasks and rest. NeuroImage, 268, 119890. 10.1016/j.neuroimage.2023.119890

Friston, K. J. (1994). Functional and effective connectivity in neuroimaging: A synthesis. Human Brain Mapping, 2(1–2), 56–78. 10.1002/hbm.460020107

Friston, K. J., Mechelli, A., Turner, R., & Price, C. J. (2000). Nonlinear Responses in fMRI: The Balloon Model, Volterra Kernels, and Other Hemodynamics. NeuroImage, 12(4), 466–477. 10.1006/nimg.2000.0630

Fukuda, M., Poplawsky, A. J., & Kim, S.-G. (2016). Submillimeter-resolution fMRI. In Progress in Brain Research (Vol. 225, pp. 123–152). Elsevier. 10.1016/bs.pbr.2016.03.003

Fukushima, M., Betzel, R. F., He, Y., De Reus, M. A., Van Den Heuvel, M. P., Zuo, X.-N., & Sporns, O. (2018). Fluctuations between high- and low-modularity topology in time-resolved functional connectivity. NeuroImage, 180, 406–416. 10.1016/j.neuroimage.2017.08.044

Fukushima, M., Betzel, R. F., He, Y., Van Den Heuvel, M. P., Zuo, X.-N., & Sporns, O. (2018). Structure–function relationships during segregated and integrated network states of human brain functional connectivity. Brain Structure and Function, 223(3), 1091–1106. 10.1007/s00429-017-1539-3

Glasser, M. F., Smith, S. M., Marcus, D. S., Andersson, J. L. R., Auerbach, E. J., Behrens, T. E. J., Coalson, T. S., Harms, M. P., Jenkinson, M., Moeller, S., Robinson, E. C., Sotiropoulos, S. N., Xu, J., Yacoub, E., Ugurbil, K., & Van Essen, D. C. (2016). The Human Connectome Project’s neuroimaging approach. Nature Neuroscience, 19(9), 1175–1187. 10.1038/nn.4361

Glasser, M. F., Sotiropoulos, S. N., Wilson, J. A., Coalson, T. S., Fischl, B., Andersson, J. L., Xu, J., Jbabdi, S., Webster, M., Polimeni, J. R., Van Essen, D. C., & Jenkinson, M. (2013). The minimal preprocessing pipelines for the Human Connectome Project. NeuroImage, 80, 105–124. 10.1016/j.neuroimage.2013.04.127

Glerean, E., Salmi, J., Lahnakoski, J. M., Jääskeläinen, I. P., & Sams, M. (2012). Functional Magnetic Resonance Imaging Phase Synchronization as a Measure of Dynamic Functional Connectivity. Brain Connectivity, 2(2), 91–101. 10.1089/brain.2011.0068

Glover, G. H., Li, T.-Q., & Ress, D. (2000). Image-based method for retrospective correction of physiological motion effects in fMRI: RETROICOR. Magnetic Resonance in Medicine, 44(1), 162–167. 10.1002/1522-2594(200007)44:1%253C162::AID-MRM23%253E3.0.CO;2-E

Gohel, S. R., & Biswal, B. B. (2015). Functional Integration Between Brain Regions at Rest Occurs in Multiple-Frequency Bands. Brain Connectivity, 5(1), 23–34. 10.1089/brain.2013.0210

Greicius, M. D., Krasnow, B., Reiss, A. L., & Menon, V. (2003). Functional connectivity in the resting brain: A network analysis of the default mode hypothesis. Proceedings of the National Academy of Sciences, 100(1), 253–258. 10.1073/pnas.0135058100

Haegens, S. (2020). Entrainment revisited: A commentary on. *Language*, Cognition and Neuroscience, 35(9), 1119–1123. 10.1080/23273798.2020.1758335

Handwerker, D. A., Roopchansingh, V., Gonzalez-Castillo, J., & Bandettini, P. A. (2012). Periodic changes in fMRI connectivity. NeuroImage, 63(3), 1712–1719. 10.1016/j.neuroimage.2012.06.078

He, B. J. (2013). Spontaneous and Task-Evoked Brain Activity Negatively Interact. The Journal of Neuroscience, 33(11), 4672–4682. 10.1523/JNEUROSCI.2922-12.2013

He, B. J., Snyder, A. Z., Zempel, J. M., Smyth, M. D., & Raichle, M. E. (2008). Electrophysiological correlates of the brain’s intrinsic large-scale functional architecture. Proceedings of the National Academy of Sciences, 105(41), 16039–16044. 10.1073/pnas.0807010105

He, B. J., Zempel, J. M., Snyder, A. Z., & Raichle, M. E. (2010). The Temporal Structures and Functional Significance of Scale-free Brain Activity. Neuron, 66(3), 353–369. 10.1016/j.neuron.2010.04.020

Honari, H., Choe, A. S., & Lindquist, M. A. (2021). Evaluating phase synchronization methods in fMRI: A comparison study and new approaches. NeuroImage, 228, 117704. 10.1016/j.neuroimage.2020.117704

Honari, H., & Lindquist, M. A. (2022). Mode decomposition-based time-varying phase synchronization for fMRI. NeuroImage, 261, 119519. 10.1016/j.neuroimage.2022.119519

Huang, N. E., Shen, Z., Long, S. R., Wu, M. C., Shih, H. H., Zheng, Q., Yen, N.-C., Tung, C. C., & Liu, H. H. (1998). The empirical mode decomposition and the Hilbert spectrum for nonlinear and non-stationary time series analysis. *Proceedings of the Royal Society of London. Series A: Mathematical*, Physical and Engineering Sciences, 454(1971), 903–995. 10.1098/rspa.1998.0193

Hutchison, R. M., Womelsdorf, T., Allen, E. A., Bandettini, P. A., Calhoun, V. D., Corbetta, M., Della Penna, S., Duyn, J. H., Glover, G. H., Gonzalez-Castillo, J., Handwerker, D. A., Keilholz, S., Kiviniemi, V., Leopold, D. A., De Pasquale, F., Sporns, O., Walter, M., & Chang, C. (2013). Dynamic functional connectivity: Promise, issues, and interpretations. NeuroImage, 80, 360–378. 10.1016/j.neuroimage.2013.05.079

Hutchison, R. M., Womelsdorf, T., Gati, J. S., Everling, S., & Menon, R. S. (2013). Resting-state networks show dynamic functional connectivity in awake humans and anesthetized macaques: Dynamic Functional Connectivity. Human Brain Mapping, 34(9), 2154–2177. 10.1002/hbm.22058

Karahanoğlu, F. I., & Van De Ville, D. (2015). Transient brain activity disentangles fMRI resting-state dynamics in terms of spatially and temporally overlapping networks. Nature Communications, 6(1), 7751. 10.1038/ncomms8751

Keilholz, S., Caballero-Gaudes, C., Bandettini, P., Deco, G., & Calhoun, V. (2017). Time-Resolved Resting-State Functional Magnetic Resonance Imaging Analysis: Current Status, Challenges, and New Directions. Brain Connectivity, 7(8), 465–481. 10.1089/brain.2017.0543

Keilholz, S. D. (2014). The Neural Basis of Time-Varying Resting-State Functional Connectivity. Brain Connectivity, 4(10), 769–779. 10.1089/brain.2014.0250

Keitel, C., Quigley, C., & Ruhnau, P. (2014). Stimulus-Driven Brain Oscillations in the Alpha Range: Entrainment of Intrinsic Rhythms or Frequency-Following Response? Journal of Neuroscience, 34(31), 10137–10140. 10.1523/JNEUROSCI.1904-14.2014

Keitel, C., Ruzzoli, M., Dugué, L., Busch, N. A., & Benwell, C. S. Y. (2022). Rhythms in cognition: The evidence revisited. European Journal of Neuroscience, 55(11–12), 2991–3009. 10.1111/ejn.15740

Khambhati, A. N., Sizemore, A. E., Betzel, R. F., & Bassett, D. S. (2018). Modeling and interpreting mesoscale network dynamics. *NeuroImage*, Brain Connectivity Dynamics, 180, 337–349. 10.1016/j.neuroimage.2017.06.029

Kwong, K. K., Belliveau, J. W., Chesler, D. A., Goldberg, I. E., Weisskoff, R. M., Poncelet, B. P., Kennedy, D. N., Hoppel, B. E., Cohen, M. S., & Turner, R. (1992). Dynamic magnetic resonance imaging of human brain activity during primary sensory stimulation. Proceedings of the National Academy of Sciences of the United States of America, 89(12), 5675–5679. 10.1073/pnas.89.12.5675

Laird, A. R., Eickhoff, S. B., Li, K., Robin, D. A., Glahn, D. C., & Fox, P. T. (2009). Investigating the Functional Heterogeneity of the Default Mode Network Using Coordinate-Based Meta-Analytic Modeling. Journal of Neuroscience, 29(46), 14496–14505. 10.1523/JNEUROSCI.4004-09.2009

Lancaster, Iatsenko, Pidde, Ticcinelli, & Stefanovska. (2018). Surrogate data for hypothesis testing in physical systems. Physics Reports, 2018(748).

Lee, H.-L., Zahneisen, B., Hugger, T., LeVan, P., & Hennig, J. (2013). Tracking dynamic resting-state networks at higher frequencies using MR-encephalography. NeuroImage, 65, 216–222. 10.1016/j.neuroimage.2012.10.015

Leonardi, N., Richiardi, J., Gschwind, M., Simioni, S., Annoni, J.-M., Schluep, M., Vuilleumier, P., & Van De Ville, D. (2013). Principal components of functional connectivity: A new approach to study dynamic brain connectivity during rest. NeuroImage, 83, 937–950. 10.1016/j.neuroimage.2013.07.019

Lindquist, M. A., Xu, Y., Nebel, M. B., & Caffo, B. S. (2014). Evaluating dynamic bivariate correlations in resting-state fMRI: A comparison study and a new approach. NeuroImage, 101, 531–546. 10.1016/j.neuroimage.2014.06.052

Liu, X., Zhang, N., Chang, C., & Duyn, J. H. (2018). Co-activation patterns in resting-state fMRI signals. NeuroImage, 180, 485–494. 10.1016/j.neuroimage.2018.01.041

Lurie, D. J., Kessler, D., Bassett, D. S., Betzel, R. F., Breakspear, M., Kheilholz, S., Kucyi, A., Liégeois, R., Lindquist, M. A., McIntosh, A. R., Poldrack, R. A., Shine, J. M., Thompson, W. H., Bielczyk, N. Z., Douw, L., Kraft, D., Miller, R. L., Muthuraman, M., Pasquini, L., … Calhoun, V. D. (2020). Questions and controversies in the study of time-varying functional connectivity in resting fMRI. Network Neuroscience, 4(1), 30–69. 10.1162/netn_a_00116

Martin, C. G., He, B. J., & Chang, C. (2021). State-related neural influences on fMRI connectivity estimation. NeuroImage, 244, 118590. 10.1016/j.neuroimage.2021.118590

Meunier, D. (2009). Hierarchical modularity in human brain functional networks. Frontiers in Neuroinformatics, 3. 10.3389/neuro.11.037.2009

Meyer, L., Sun, Y., & Martin, A. E. (2020). Synchronous, but not entrained: Exogenous and endogenous cortical rhythms of speech and language processing. Language, Cognition and Neuroscience, 35(9), 1089–1099. 10.1080/23273798.2019.1693050

Miller, R. L., Abrol, A., Adali, T., Levin-Schwarz, Y., & Calhoun, V. D. (2018). Resting-State fMRI Dynamics and Null Models: Perspectives, Sampling Variability, and Simulations. Frontiers in Neuroscience, 12, 551. 10.3389/fnins.2018.00551

Mitra, A., Snyder, A. Z., Hacker, C. D., & Raichle, M. E. (2014). Lag structure in resting-state fMRI. Journal of Neurophysiology, 111(11), 2374–2391. 10.1152/jn.00804.2013

Mucha, P. J., Richardson, T., Macon, K., Porter, M. A., & Onnela, J.-P. (2010). Community Structure in Time-Dependent, Multiscale, and Multiplex Networks. Science, 328(5980), 876–878. 10.1126/science.1184819

Niazy, R. K., Xie, J., Miller, K., Beckmann, C. F., & Smith, S. M. (2011). Spectral characteristics of resting state networks. In Progress in Brain Research (Vol. 193, pp. 259–276). Elsevier. 10.1016/B978-0-444-53839-0.00017-X

Norcia, A. M., Appelbaum, L. G., Ales, J. M., Cottereau, B. R., & Rossion, B. (2015). The steady-state visual evoked potential in vision research: A review. Journal of Vision, 15(6), 4. 10.1167/15.6.4

Ogawa, S., Lee, T. M., Kay, A. R., & Tank, D. W. (1990). Brain magnetic resonance imaging with contrast dependent on blood oxygenation. Proceedings of the National Academy of Sciences of the United States of America, 87(24), 9868–9872. 10.1073/pnas.87.24.9868

Pedersen, M., Omidvarnia, A., Zalesky, A., & Jackson, G. D. (2018). On the relationship between instantaneous phase synchrony and correlation-based sliding windows for time-resolved fMRI connectivity analysis. NeuroImage, 181, 85–94. 10.1016/j.neuroimage.2018.06.020

Penttonen, M. (2003). Natural logarithmic relationship between brain oscillators. Thalamus & Related Systems, 2(2), 145–152. 10.1016/S1472-9288(03)00007-4

Ponce-Alvarez, A., Deco, G., Hagmann, P., Romani, G. L., Mantini, D., & Corbetta, M. (2015). Resting-State Temporal Synchronization Networks Emerge from Connectivity Topology and Heterogeneity. PLOS Computational Biology, 11(2), e1004100. 10.1371/journal.pcbi.1004100

Preti, M. G., Bolton, T. A., & Van De Ville, D. (2017). The dynamic functional connectome: State-of-the-art and perspectives. NeuroImage, 160, 41–54. 10.1016/j.neuroimage.2016.12.061

Preti, M. G., Haller, S., Giannakopoulos, P., & Van De Ville, D. (2015). Decomposing dynamic functional connectivity onto phase-dependent eigenconnectivities using the Hilbert transform. 2015 IEEE 12th International Symposium on Biomedical Imaging (ISBI), 38–41. 10.1109/ISBI.2015.7163811

Quigley, C. (2022). Forgotten rhythms? Revisiting the first evidence for rhythms in cognition. The European Journal of Neuroscience, 55(11–12), 3266–3276. 10.1111/ejn.15450

Raichle, M. E., MacLeod, A. M., Snyder, A. Z., Powers, W. J., Gusnard, D. A., & Shulman, G. L. (2001). A default mode of brain function. Proceedings of the National Academy of Sciences, 98(2), 676–682. 10.1073/pnas.98.2.676

Reid, A. T., Headley, D. B., Mill, R. D., Sanchez-Romero, R., Uddin, L. Q., Marinazzo, D., Lurie, D. J., Valdés-Sosa, P. A., Hanson, S. J., Biswal, B. B., Calhoun, V., Poldrack, R. A., & Cole, M. W. (2019). Advancing functional connectivity research from association to causation. Nature Neuroscience, 22(11), 1751–1760. 10.1038/s41593-019-0510-4

Sakoğlu, Ü., Pearlson, G. D., Kiehl, K. A., Wang, Y. M., Michael, A. M., & Calhoun, V. D. (2010). A method for evaluating dynamic functional network connectivity and task-modulation: Application to schizophrenia. *Magnetic Resonance Materials in Physics*, Biology and Medicine, 23(5–6), 351–366. 10.1007/s10334-010-0197-8

Salvador, R., Martínez, A., Pomarol-Clotet, E., Gomar, J., Vila, F., Sarró, S., Capdevila, A., & Bullmore, E. (2008). A simple view of the brain through a frequency-specific functional connectivity measure. NeuroImage, 39(1), 279–289. 10.1016/j.neuroimage.2007.08.018

Schaefer, A., Kong, R., Gordon, E. M., Laumann, T. O., Zuo, X.-N., Holmes, A. J., Eickhoff, S. B., & Yeo, B. T. T. (2018). Local-Global Parcellation of the Human Cerebral Cortex from Intrinsic Functional Connectivity MRI. Cerebral Cortex, 28(9), 3095–3114. 10.1093/cercor/bhx179

Shine, J. M., Bissett, P. G., Bell, P. T., Koyejo, O., Balsters, J. H., Gorgolewski, K. J., Moodie, C. A., & Poldrack, R. A. (2016). The Dynamics of Functional Brain Networks: Integrated Network States during Cognitive Task Performance. Neuron, 92(2), 544–554. 10.1016/j.neuron.2016.09.018

Smith, S. M., Fox, P. T., Miller, K. L., Glahn, D. C., Fox, P. M., Mackay, C. E., Filippini, N., Watkins, K. E., Toro, R., Laird, A. R., & Beckmann, C. F. (2009). Correspondence of the brain’s functional architecture during activation and rest. Proceedings of the National Academy of Sciences, 106(31), 13040–13045. 10.1073/pnas.0905267106

Sporns, O. (2013). Network attributes for segregation and integration in the human brain. Current Opinion in Neurobiology, 23(2), 162–171. 10.1016/j.conb.2012.11.015

Sporns, O., Tononi, G., & Edelman, G. M. (2000). Connectivity and complexity: The relationship between neuroanatomy and brain dynamics. Neural Networks, 13(8–9), 909–922. 10.1016/S0893-6080(00)00053-8

Strindberg, M., Fransson, P., Cabral, J., & Ådén, U. (2021). Spatiotemporally flexible subnetworks reveal the quasi-cyclic nature of integration and segregation in the human brain. NeuroImage, 239, 118287. 10.1016/j.neuroimage.2021.118287

Tagliazucchi, E., Von Wegner, F., Morzelewski, A., Brodbeck, V., & Laufs, H. (2012). Dynamic BOLD functional connectivity in humans and its electrophysiological correlates. Frontiers in Human Neuroscience, 6. 10.3389/fnhum.2012.00339

Thompson, W. H., & Fransson, P. (2015). The frequency dimension of fMRI dynamic connectivity: Network connectivity, functional hubs and integration in the resting brain. NeuroImage, 121, 227–242. 10.1016/j.neuroimage.2015.07.022

Thompson, W. H., & Fransson, P. (2018). A common framework for the problem of deriving estimates of dynamic functional brain connectivity. NeuroImage, 172, 896–902. 10.1016/j.neuroimage.2017.12.057

Thompson, W. H., Richter, C. G., Plavén-Sigray, P., & Fransson, P. (2018). Simulations to benchmark time-varying connectivity methods for fMRI. PLOS Computational Biology, 14(5), e1006196. 10.1371/journal.pcbi.1006196

Torres, M. E., Colominas, M. A., Schlotthauer, G., & Flandrin, P. (2011). A complete ensemble empirical mode decomposition with adaptive noise. 2011 IEEE International Conference on Acoustics, Speech and Signal Processing (ICASSP), 4144–4147. 10.1109/ICASSP.2011.5947265

Uddin, L. Q., Betzel, R. F., Cohen, J. R., Damoiseaux, J. S., De Brigard, F., Eickhoff, S. B., Fornito, A., Gratton, C., Gordon, E. M., Laird, A. R., Larson-Prior, L., McIntosh, A. R., Nickerson, L. D., Pessoa, L., Pinho, A. L., Poldrack, R. A., Razi, A., Sadaghiani, S., Shine, J. M., … Spreng, R. N. (2023). Controversies and progress on standardization of large-scale brain network nomenclature. Network Neuroscience, 7(3), 864–905. 10.1162/netn_a_00323

Uddin, L. Q., Clare Kelly, A. M., Biswal, B. B., Xavier Castellanos, F., & Milham, M. P. (2009). Functional connectivity of default mode network components: Correlation, anticorrelation, and causality. Human Brain Mapping, 30(2), 625–637. 10.1002/hbm.20531

Uğurbil, K., Xu, J., Auerbach, E. J., Moeller, S., Vu, A. T., Duarte-Carvajalino, J. M., Lenglet, C., Wu, X., Schmitter, S., Van De Moortele, P. F., Strupp, J., Sapiro, G., De Martino, F., Wang, D., Harel, N., Garwood, M., Chen, L., Feinberg, D. A., Smith, S. M., … Yacoub, E. (2013). Pushing spatial and temporal resolution for functional and diffusion MRI in the Human Connectome Project. NeuroImage, 80, 80–104. 10.1016/j.neuroimage.2013.05.012

Van De Ville, D., & Liégeois, R. (2024). Dynamic functional connectivity to tile the spatiotemporal mosaic of brain states. Imaging Neuroscience, 2, imag–2–00364. 10.1162/imag_a_00364

Van Essen, D. C., Smith, S. M., Barch, D. M., Behrens, T. E. J., Yacoub, E., & Ugurbil, K. (2013). The WU-Minn Human Connectome Project: An overview. *NeuroImage*, Mapping the Connectome, 80, 62–79. 10.1016/j.neuroimage.2013.05.041

Varela, F., Lachaux, J.-P., Rodriguez, E., & Martinerie, J. (2001). The brainweb: Phase synchronization and large-scale integration. Nature Reviews Neuroscience, 2(4), 229–239. 10.1038/35067550

Vidaurre, D. (2024). Dynamic functional connectivity: Why the controversy? Imaging Neuroscience, 2, imag–2–00363. 10.1162/imag_a_00363

Vidaurre, D., Smith, S. M., & Woolrich, M. W. (2017). Brain network dynamics are hierarchically organized in time. Proceedings of the National Academy of Sciences, 114(48), 12827–12832. 10.1073/pnas.1705120114

Wong, K.-F., & Wang, X.-J. (2006). A Recurrent Network Mechanism of Time Integration in Perceptual Decisions. The Journal of Neuroscience, 26(4), 1314–1328. 10.1523/JNEUROSCI.3733-05.2006

Xu, B., Sheng, Y., Li, P., Cheng, Q., & Wu, J. (2019). Causes and classification of EMD mode mixing. Vibroengineering Procedia, 22, 158–164. 10.21595/vp.2018.20250

Yaesoubi, M., Allen, E. A., Miller, R. L., & Calhoun, V. D. (2015). Dynamic coherence analysis of resting fMRI data to jointly capture state-based phase, frequency, and time-domain information. NeuroImage, 120, 133–142. 10.1016/j.neuroimage.2015.07.002

Yeo, B. T. T., Krienen, F. M., EickhoT, S. B., Yaakub, S. N., Fox, P. T., Buckner, R. L., Asplund, C. L., & Chee, M. W. L. (2015). Functional Specialization and Flexibility in Human Association Cortex. Cerebral Cortex, 25(10), 3654–3672. 10.1093/cercor/bhu217

Yeo, B. T. T., Krienen, F. M., Sepulcre, J., Sabuncu, M. R., Lashkari, D., Hollinshead, M., Roffman, J. L., Smoller, J. W., Zöllei, L., Polimeni, J. R., Fischl, B., Liu, H., & Buckner, R. L. (2011). The organization of the human cerebral cortex estimated by intrinsic functional connectivity. Journal of Neurophysiology, 106(3), 1125–1165. 10.1152/jn.00338.2011

Zalesky, A., Fornito, A., Cocchi, L., Gollo, L. L., & Breakspear, M. (2014). Time-resolved resting-state brain networks. Proceedings of the National Academy of Sciences, 111(28), 10341–10346. 10.1073/pnas.1400181111

Zatman, M. (1997). How narrow is narrowband? [Adaptive array signal processing]. Conference Record of the Thirty-First Asilomar Conference on Signals, Systems and Computers (Cat. No.97CB36136), 2, 1341–1345 vol.2. 10.1109/ACSSC.1997.679122

Zuo, X.-N., Di Martino, A., Kelly, C., Shehzad, Z. E., Gee, D. G., Klein, D. F., Castellanos, F. X., Biswal, B. B., & Milham, M. P. (2010). The oscillating brain: Complex and reliable. NeuroImage, 49(2), 1432–1445. 10.1016/j.neuroimage.2009.09.037

