## Supplementary information for "Time-resolved brain network community detection based on instantaneous phase of fMRI data"

### Supplementary Materials, Strindberg et al., 2026

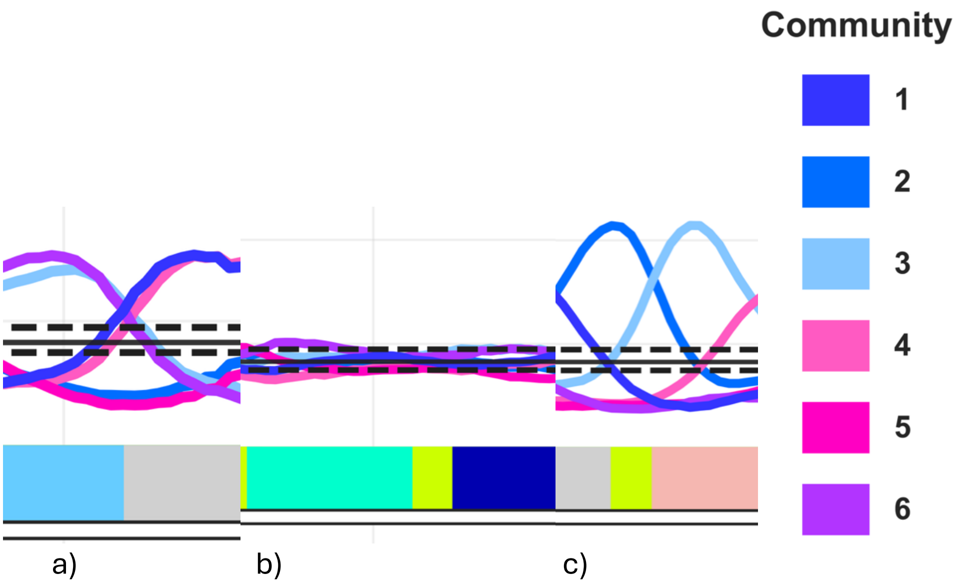


**Fig. S1.** Three different types of states related to integration/segregation between phase communities. (a) “bimodal segregation”, (b) “equalized”, and (c) “single community dominating”. In the bimodal segregated state, two communities that are maximally segregated (in terms of instantaneous phase) have each an almost equal number of parcels assigned to them on average (across subjects). In the equalized state, all communities are similar in parcel size and hence close to or within the non-significant statistical boundaries. In the single dominating community case, one community is substantially larger than the others. Excerpts of data are taken from the fMRI motor task.

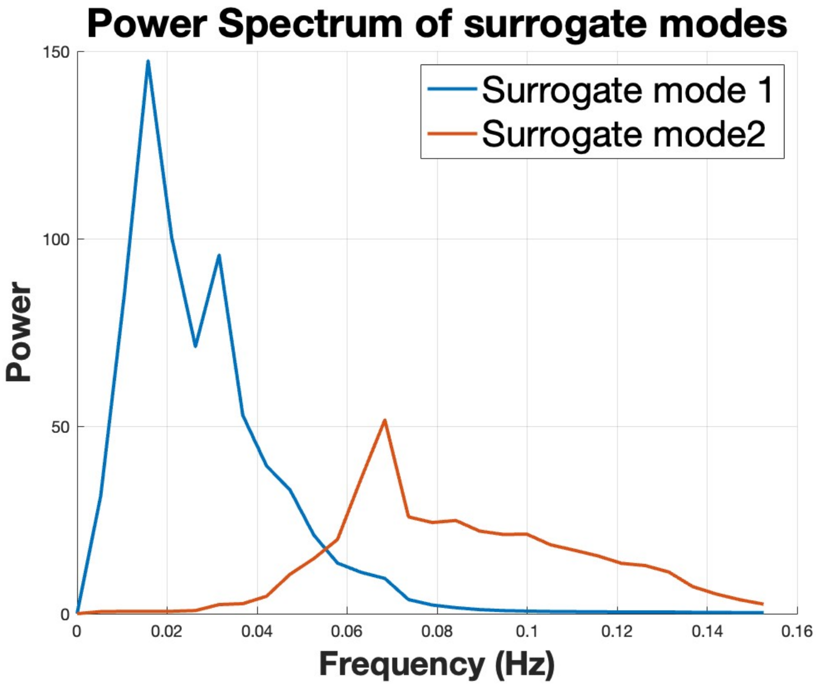


**Fig. S2.** Power spectrum for phase-scrambled surrogate data.

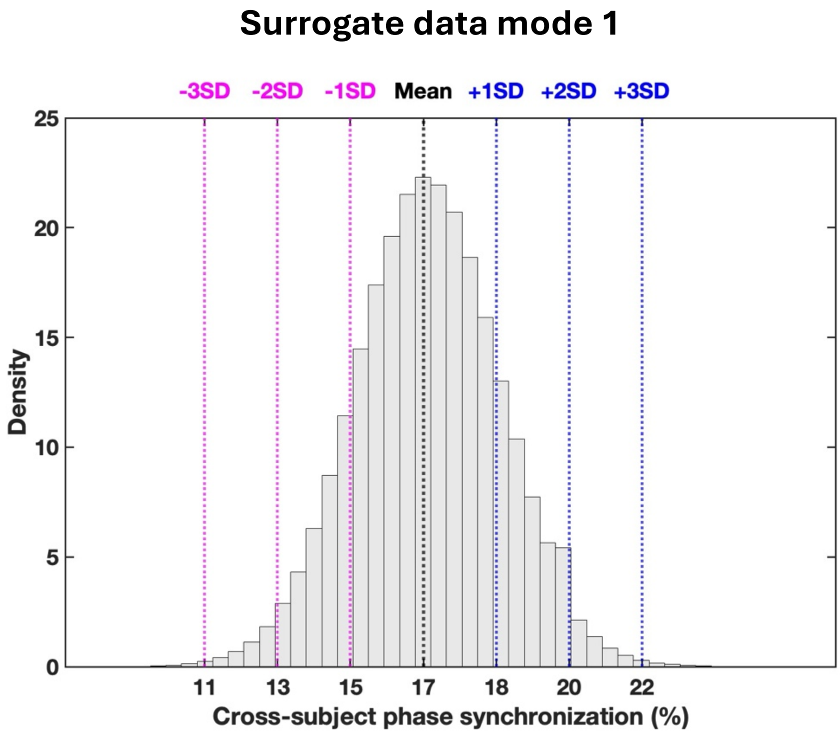


a.

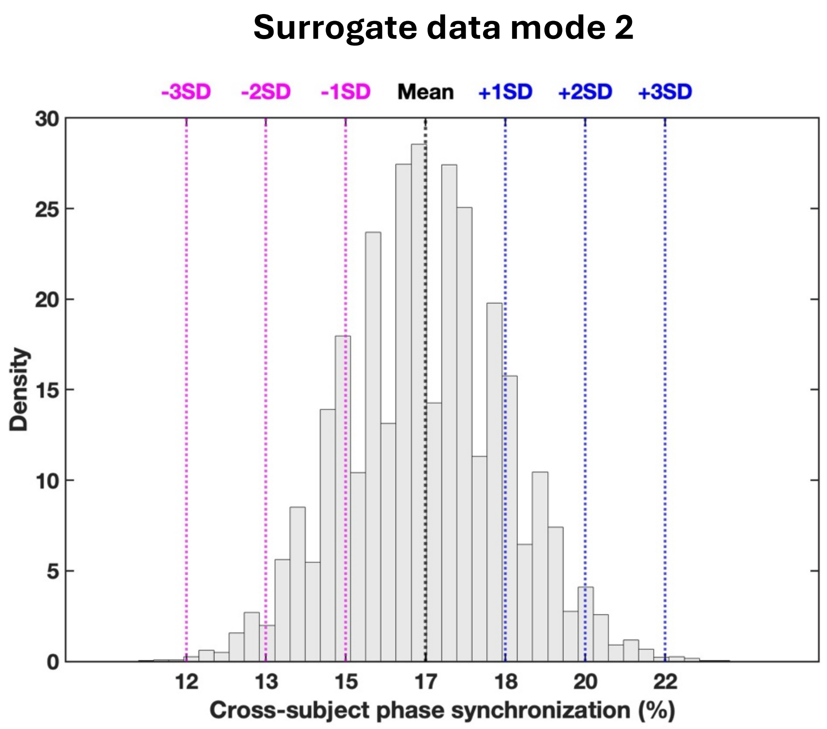


b.

**Fig. S3.** Distribution of cross-subject phase synchronization for surrogate data. Aggregation of all data across all subjects, time-points and six communities. The synchronization data is normally distributed and the interval -3SD to +3SD encompasses 99.7% of the data. (a) mode 1 (lower frequency range), max = 0.27, min = 0.08, mean = 0.167, -3SD = 0.11, +3SD = 0.22. (b) mode 2 (higher frequency range), max = 0.25, min = 0.09, mean = 0.167, -3SD = 0.12, +3SD = 0.22.

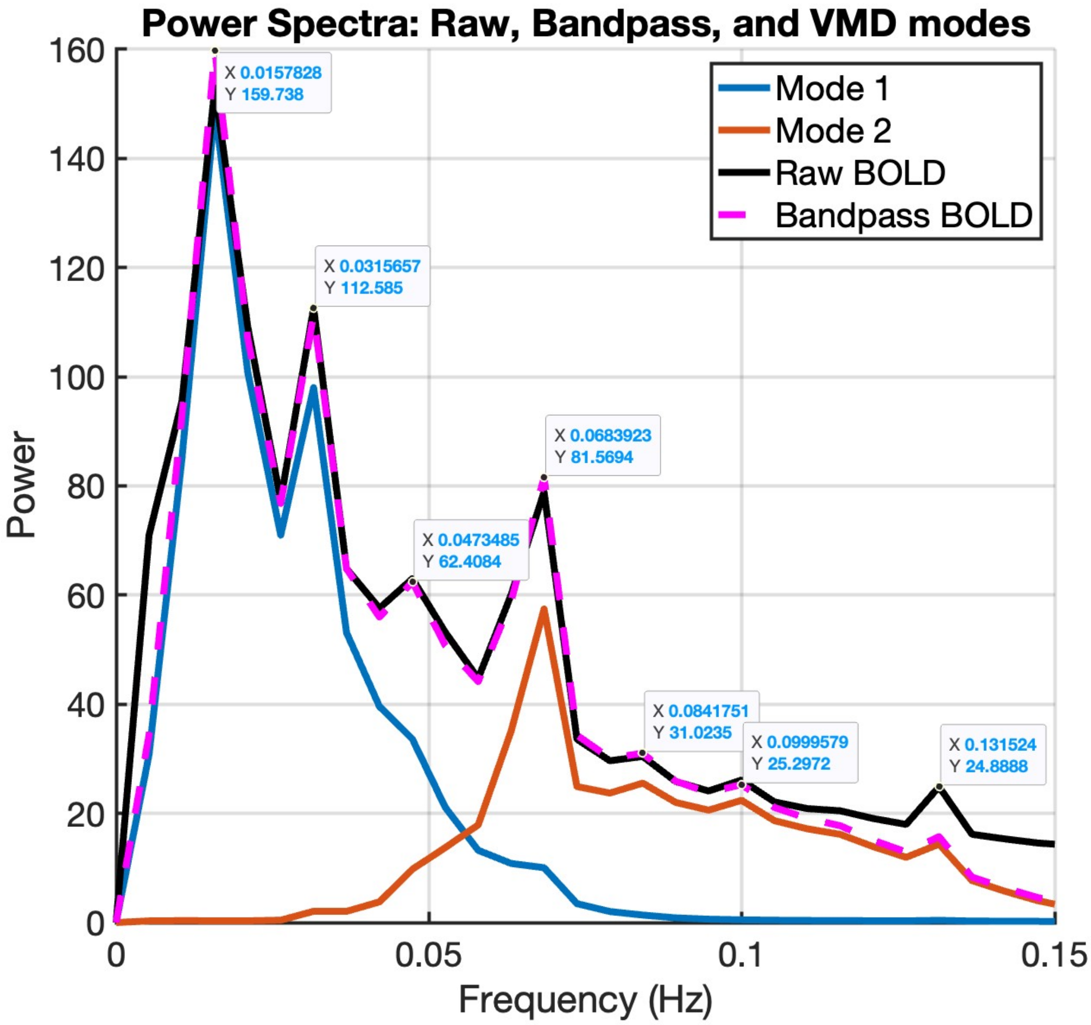


**Fig. S4.** Power spectrum for mode 1 (blue), mode 2 (red), bandpass filtered fMRI motor data (dotted magenta) and raw (pre-processed) BOLD data (black line).

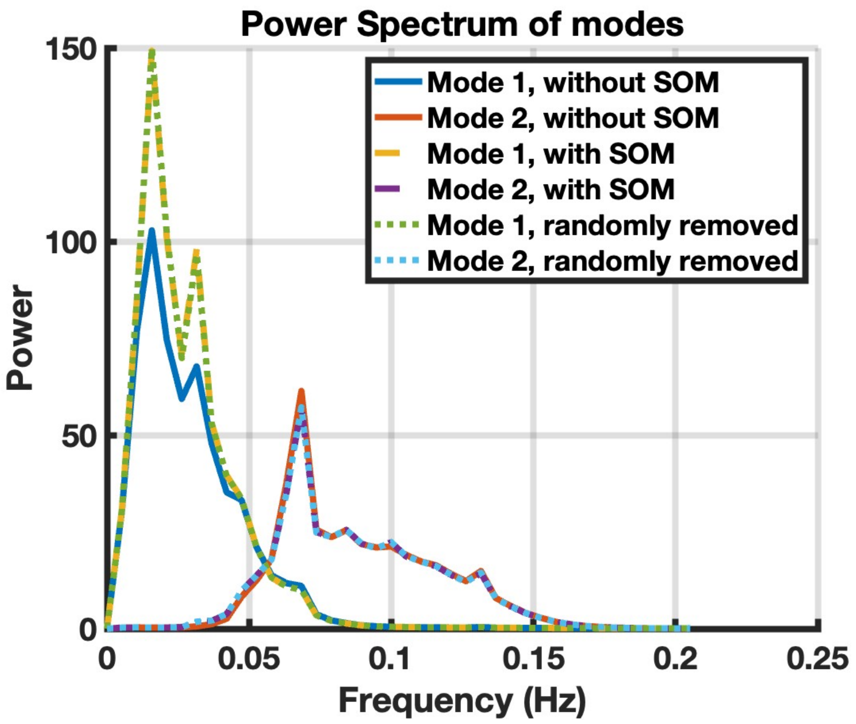


**Fig. S5a.** Power spectrum with SOM parcels removed compared to equal number of randomly removed parcels (n=77) and all parcels. SOM parcels make a substantial contribution to both peaks in mode 1.

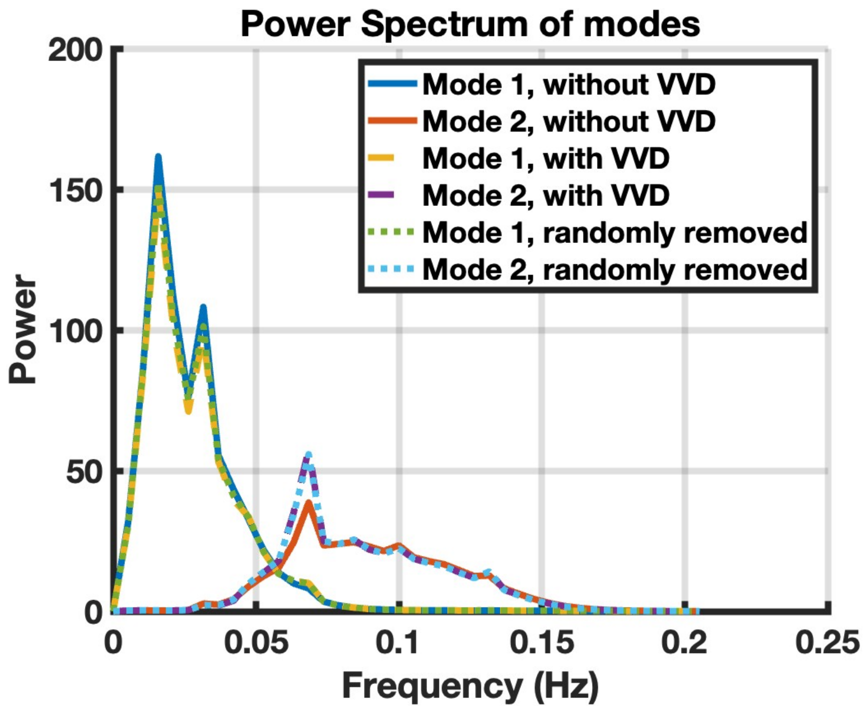

**Fig. S5b.** Power spectrum with VVD (VIS, VAN, DAN) parcels removed compared to equal number of randomly removed parcels (n=154) and all parcels. VVD parcels make a substantial contribution to the main peak in mode 2.

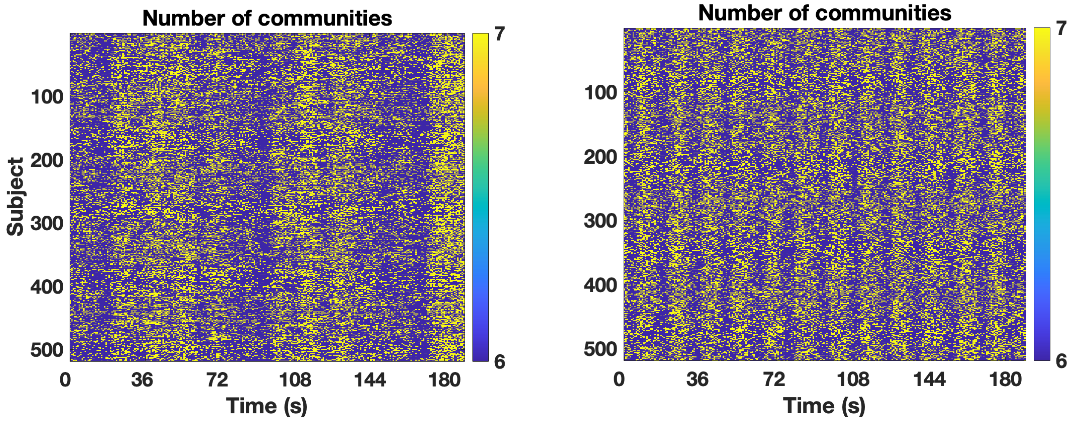


a. Mode 1 b. Mode 2

**Fig. S6**. Number of unique communities as function of time. The appearance of an additional seventh community was seen both during task and rest periods (see also Fig. 2) with a normalisation of average number of communities to baseline (n=6) during cue presentation. The effect was more pronounced in the higher frequencies range (mode 2). A potential interpretation is that cue presentation is associated with heightened attention which would be associated with more integration (i.e. baseline number of communities).

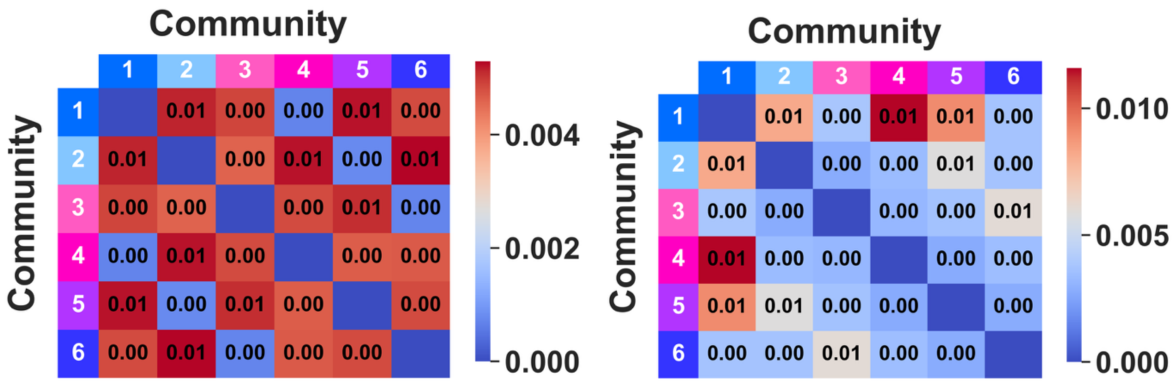


a) Average SD cosine b) Average SD sine

**Fig. S7.** Average standard deviation of average difference of instantaneous phase between communities a) cosine, b) sine.

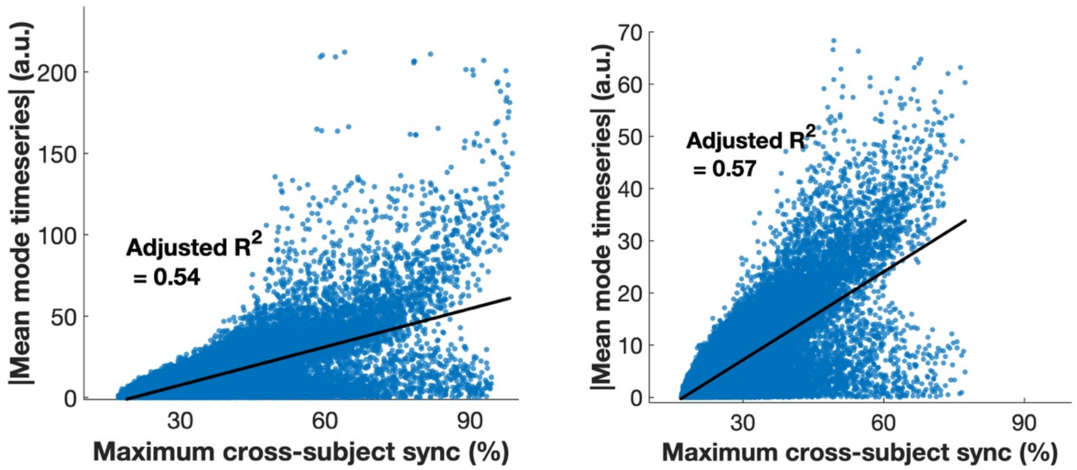


**Fig. S8.** Relationship between maximum (across 6 communities) cross-subject phase synchronization and amplitude of the mean (across subjects) mode timeseries. (a) mode 1. (b) mode 2. The relationship between absolute value of the mean mode amplitude and maximum cross-subject phase synchronization for all timepoints are plotted in supplement Fig. S8. Of note, strong synchronization was seen at amplitudes close to zero indicating zero-crossings where the amplitudes transition between positive and negative values. It was also seen for negative amplitudes (here shifted to positives to calculate the statistics)

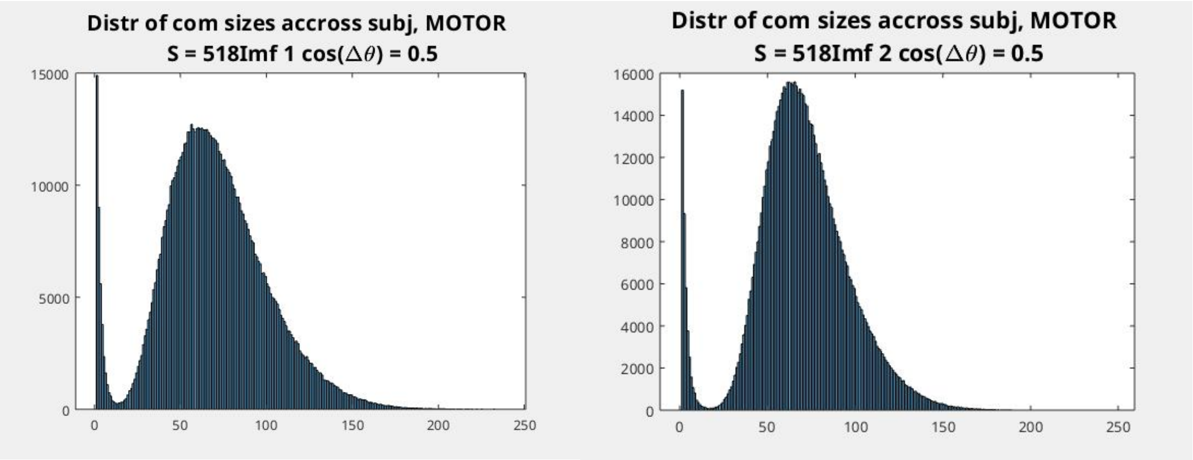


**a). Mode 1** **b) Mode 2**

**Fig. S9.** Distribution of community size (number of parcels) for mode 1 (a) and mode 2 (b). All subjects, timepoints and communities are aggregated.

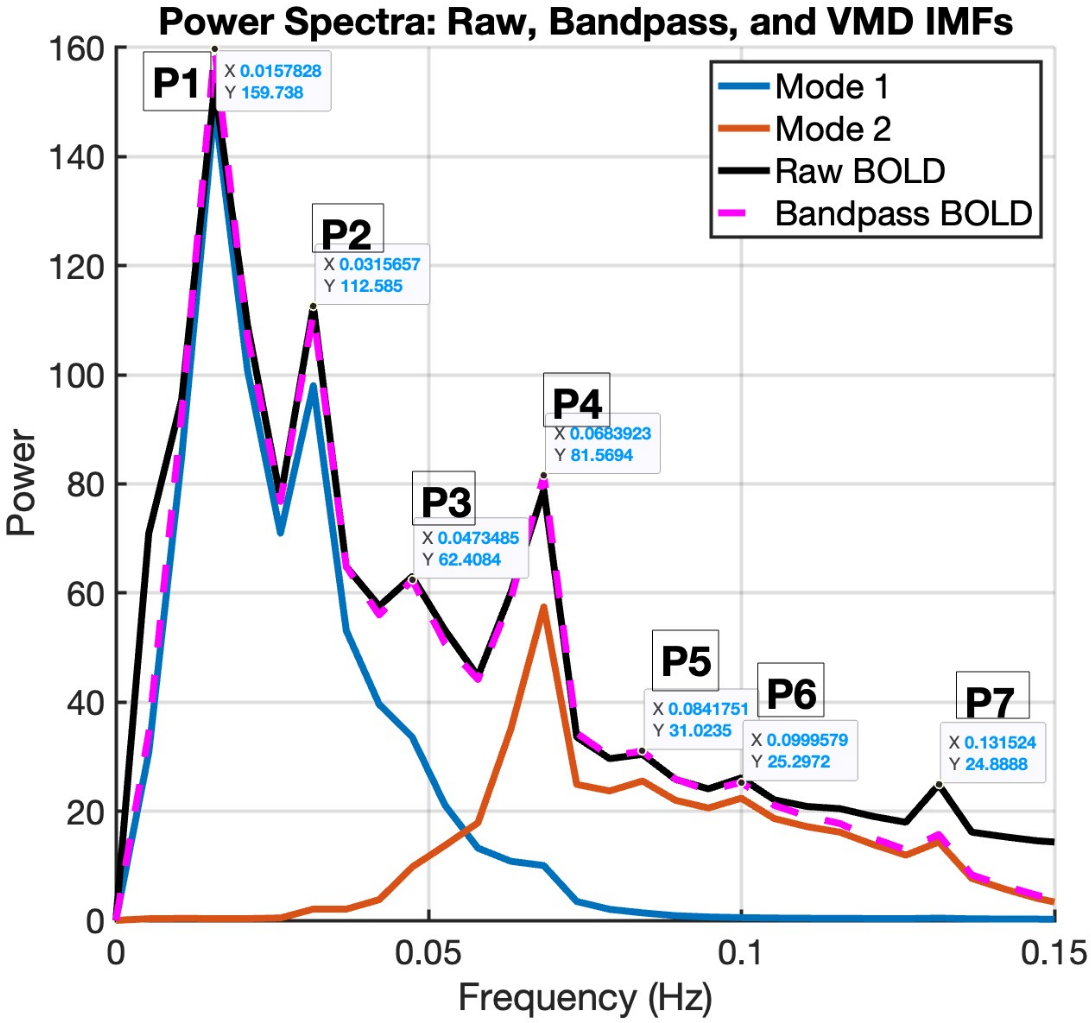

**Fig. S10.** Higher order harmonics in the power-spectra for the fMRI motor data. Harmonics of the first (and largest) peak (P1) at 0.016 Hz are seen at 0.032 Hz (2*P1, peak 2), 0.047Hz (3*P1, peak 3). While close, P4-P6 are not true harmonics.  P7 is most likely aliasing of cardiac rhythms which could also be the case for P3, P5 and P6.
